# From clusters to curves: A model-based alternative to cluster-based permutation tests for the analysis of time-resolved data

**DOI:** 10.1101/2025.08.29.672336

**Authors:** Ladislas Nalborczyk, Paul Bürkner

**Affiliations:** Aix Marseille Univ, CNRS, LPL; TU Dortmund University, Department of Statistics

**Keywords:** EEG, MEG, cluster-based permutation tests, generalised additive models, mixed-effects models, multilevel models, Bayesian statistics, brms

## Abstract

Time-resolved electrophysiological measurements such as those obtained through magneto- and electroencephalography (M/EEG) offer a unique window onto the neural activity underlying cognitive processes. Researchers are often interested in determining whether and when these signals differ across experimental conditions or participant groups. The conventional approach involves mass univariate statistical testing across time and space followed by corrections for multiple comparisons or some form of cluster-based inference. While effective for controlling error rates at the cluster-level, cluster-based inference comes with a significant limitation: by shifting the focus of inference from individual time points to clusters, it prevents drawing conclusions about the precise onset or offset of observed effects. Here, we present a *model-based* alternative for analysing M/EEG time series, such as event-related potentials or time-resolved decoding accuracy. Our approach leverages Bayesian generalised additive multilevel models, providing posterior odds at each time point, while naturally accounting for between-participant variability. Using simulated data with known ground truth, we show that the proposed approach consistently provides accurate recovery of the overall temporal extent and onset of neural effects across a range of signal-to-noise ratios, although offset localisation is comparatively more variable. In an empirical MEG decoding dataset, the approach also yields the highest split-half temporal overlap among the methods considered and highly stable temporal segmentation across repeated participant splits. Together, these results illustrate the potential of model-based inference as a complementary approach to conventional cluster-based procedures when temporally localised inference and explicit modelling of between-participant variability are scientifically important. We provide an open-source R package implementing the method and describe how it can be integrated into M/EEG analysis pipelines using MNE-Python.

## 1 Introduction

### 1.1 Problem statement

Understanding the temporal dynamics of cognitive processes requires methods that can capture fast-changing neural activity with high temporal resolution. Magnetoencephalography and electroencephalography (M/EEG) are two such methods, widely used in cognitive neuroscience for their ability to track brain activity at the millisecond scale. These techniques provide rich time series data that reflect how neural responses unfold in response to stimuli or tasks. A central goal in many M/EEG studies is to determine whether, when, and where neural responses differ across experimental conditions or groups.

The conventional approach for addressing such questions relies on mass-univariate statistical testing across time and/or sensors, followed by corrections for multiple comparisons to maintain nominal error rates (e.g., FWER or FDR). Cluster-based permutation inference is the most common strategy in M/EEG research and is implemented as a recommended workflow in widely used software ecosystems (e.g., EEGLAB, Delorme & Makeig, 2004; MNE-Python, Gramfort, 2013). While effective for controlling error rates at the cluster level, cluster-based permutation tests also impose important constraints on what can be concluded. By design, they shift inference away from individual samples (time points, sensors, voxels) toward clusters formed from contiguous supra-threshold test statistics, which limits the interpretability of spatial and temporal localisation (Maris & Oostenveld, 2007; Sassenhagen & Draschkow, 2019). Importantly, the limitations of cluster-based inference extend beyond temporal imprecision in onset/offset estimation. Cluster-based outcomes can be sensitive to analysis choices (e.g., cluster-forming threshold or TFCE settings, significance threshold, test statistic, and crucially, the analysis window) and do not provide an explicit probabilistic model for how effects evolve over time at the population level.

To address these limitations, we introduce a model-based framework for time-resolved M/EEG outcomes based on Bayesian generalised additive multilevel models. The core idea is to model the expected signal over time as a smooth function, with hierarchical structure to represent between-participant variability, and to quantify evidence for above-baseline (or above-chance) effects directly in the posterior distribution at the level of individual time points (and, by extension, sensors/voxels). This approach regularises the mean time course via smoothness priors and yields coherent uncertainty quantification for time-localised quantities. Using simulated data with known ground truth and an empirical MEG decoding dataset, we show how this framework complements standard permutation-based approaches: it provides interpretable, model-based uncertainty about when effects emerge and subside, while making the role of modelling choices (e.g., basis dimension) explicit and amenable to sensitivity analysis.

### 1.2 Statistical errors and cluster-based inference

The issues with multiple comparisons represent a common and well-recognised danger in neuroimaging and M/EEG research, where the collected data allow for a multitude of potential hypothesis tests and is characterised by complex structures of spatiotemporal dependencies. The probability of obtaining at least one false positive in an ensemble (family) of *m* independent tests (i.e., the FWER) is computed as 1 − (1 *− α*)*^m^* (for *m* = 10 independent tests and *α* = 0.05, it is approximately equal to 0.4). Different methods exist to control the FWER, that is, to bring it back to *α*. Most methods apply a simple correction to series of *p*-values issued from univariate statistical tests (e.g., t-tests). For instance, the Bonferroni correction sets the per-test significance threshold to *α*/*m*, or equivalently multiplies each *p*-value by *m*. This procedure controls the FWER without requiring independence among tests, but can be conservative because it does not exploit their dependence structure. This is particularly relevant for M/EEG data, in which neighbouring temporal and spatial samples are often strongly correlated. Alternative procedures control the false discovery rate (FDR), defined as the expected proportion of false discoveries among rejected hypotheses (e.g., Benjamini & Hochberg, 1995; Benjamini & Yekutieli, 2001). Although these procedures provide useful multiplicity control, they do not explicitly model the spatial and temporal structure of M/EEG signals.

A popular technique to account for spatiotemporal dependencies while controlling the FWER is cluster-based inference (Bullmore et al., 1999; Maris & Oostenveld, 2007). A typical cluster-based permutation procedure involves two successive steps (for more details on cluster-based inference, see for instance Frossard & Renaud, 2022; Maris, 2011; Maris & Oostenveld, 2007; Sassenhagen & Draschkow, 2019). First, clusters are defined as sets of contiguous time points, sensors, voxels, etc, whose activity, summarised by some test statistic (e.g., a *t*-value), exceeds a predefined threshold (e.g., the 95th percentile of the parametric null distribution). Clusters are then characterised by their height (i.e., maximal value), extent (number of constituent elements), or some combination of both, for instance by summing the statistics within a cluster, an approach referred to as “cluster mass” (Maris & Oostenveld, 2007; Pernet et al., 2015). Then, the null hypothesis is tested by computing a *p*-value for each identified cluster by comparing its mass with the null distribution of cluster masses (obtained via permutation). A significant cluster indicates that the cluster-level statistic is unlikely under the null hypothesis; it does not imply that each constituent sample is individually significant. As such, it would be incorrect to conclude, for instance, that the time point of a significant cluster is the first moment at which some conditions differ (Frossard & Renaud, 2022; Sassenhagen & Draschkow, 2019). Because the inference is performed at the second step (i.e., once clusters have been formed), no conclusion can be made about individual datapoints (e.g., time points, sensors, etc). More recent procedures such as ClusterDepth provide strong FWER-controlled inference at the level of individual samples while retaining a cluster-based framework, although their performance for precise onset localisation can remain limited.

As different cluster-forming thresholds lead to clusters with different spatial or temporal extent, this initial threshold modulates the sensitivity of the subsequent permutation test. The threshold-free cluster enhancement (TFCE) method was introduced by S. Smith & Nichols (2009) to avoid the need to select a single arbitrary cluster-forming threshold. Rather than forming clusters at one threshold, TFCE evaluates each sample across a range of thresholds. At a given threshold *h*, the contribution to the TFCE score depends jointly on the height of the threshold and on the extent *e_t_*(*h*) of the cluster containing sample *t*. The TFCE score at time point *t* can be written as:

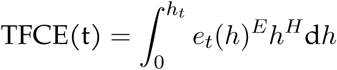

where *h_t_* is the observed statistic at time point *t*, and the parameters *E* and *H* control the relative contributions of cluster extent and height, respectively (typically *E* = 0.5 and *H* = 2). In practice, this integral is approximated numerically over a series of thresholds. Statistical significance is then assessed by permutation: for each permutation, the maximum TFCE statistic across the analysed samples is retained to construct a null distribution, against which the observed sample-wise TFCE scores are compared. This procedure can therefore provide FWER-controlled inference at the sample level without requiring a single cluster-forming threshold (e.g., Pernet et al., 2015; S. Smith & Nichols, 2009). Nevertheless, simulation studies have shown that the temporal boundaries identified by cluster-based procedures, including TFCE, can provide biased or imprecise estimates of effect onset and offset (Rousselet, 2025; Sassenhagen & Draschkow, 2019).

### 1.3 Further limitations of cluster-based permutation tests

Although cluster-based permutation tests are widely used and often appropriate for controlling error rates under a global null, they come with several limitations that are especially relevant for time-resolved M/EEG questions. First, temporal dependencies are not explicitly modelled. Cluster-based inference does not specify a generative model for temporal dependence in the measured signal. Instead, temporal structure is summarised post-hoc by identifying clusters of adjacent supra-threshold test statistics (e.g., t- or F-values). Second, within-participant variability is typically not explicitly modelled. In many workflows, data are averaged within participants (e.g., ERPs, decoding time courses) prior to group inference. Although this reduces data dimensionality, it discards information about within-participant variability and differences in measurement precision across participants. Consequently, participant-level estimates are typically treated as equally reliable, potentially reducing statistical efficiency and limiting the explicit modelling of different sources of uncertainty. Third, participant-level clusters (when extracted) do not yield a principled population model of timing variability. It is possible to extract clusters per participant, but the resulting onset/offset summaries do not by themselves quantify between-participant variability unless an explicit second-stage model is specified. Fourth, results can vary substantially with several analysis “hyper-parameters”. These include the cluster-forming threshold (or TFCE parameters), the significance threshold and tail definition, the choice of test statistic, the temporal/spatial analysis window, and pre-processing decisions that affect temporal resolution (e.g., filtering, decimation).

Together, these considerations motivate a complementary approach in which the temporal trajectory is itself part of the statistical model, uncertainty can be propagated to time-localised quantities, and between-participant variability can be represented explicitly. This idea is not entirely new: regression-based and smooth-model approaches have increasingly been used to estimate electrophysiological responses. We therefore first review this work before introducing the GAM framework used here.

### 1.4 Previous work on modelling M/EEG data

M/EEG signals capture neural activity originating from various brain regions and are often contaminated by artefacts unrelated to the cognitive processes under investigation. Consequently, analysing M/EEG data necessitates methods that can disentangle task-relevant neural signals from extraneous “noise.” A widely adopted technique for this purpose is the estimation of event-related potentials (ERPs), which are stereotyped electrophysiological responses time-locked to specific sensory, cognitive, or motor events. Typically, ERPs are derived by averaging EEG or MEG epochs across multiple trials aligned to the event of interest (e.g., stimulus onset), thereby enhancing the signal-to-noise ratio by attenuating non-time-locked activity. However, this averaging approach has notable limitations: it assumes consistent latency and amplitude across trials and is primarily suited for simple categorical designs. Such assumptions may not hold in more complex experimental paradigms, potentially leading to suboptimal ERP estimations (e.g., N. J. Smith & Kutas, 2014a).

To overcome the limitations of simple averaging, several model-based approaches for estimating ERPs have been proposed. These methods are motivated by the observation that traditional ERP averaging is mathematically equivalent to fitting an intercept-only linear regression model in a simple categorical design without overlapping events (N. J. Smith & Kutas, 2014a). In contrast to simple averaging, regression-based approaches to ERP estimation offer substantially greater flexibility. Notably, they allow for the modelling of both linear and nonlinear effects of continuous predictors, such as word frequency or age (e.g., N. J. Smith & Kutas, 2014a, 2014b; Tremblay & Newman, 2014), and enable the disentangling of overlapping cognitive processes (e.g., Ehinger & Dimigen, 2019; Skukies et al., 2024; Skukies & Ehinger, 2021). One widely used implementation of this approach is provided by the LIMO EEG toolbox (Pernet et al., 2011), which follows a multi-stage analysis pipeline. First, a separate regression model is fit for each datapoint (e.g., each time point and electrode) at the individual level to estimate ERP responses. This is followed by group-level statistical analyses of the resulting regression coefficients, often accompanied by corrections for multiple comparisons or cluster-based inference (for recent applied examples, see Dunagan et al., 2025; Wüllhorst et al., 2025).

Although this framework allows for the inclusion of a wide range of predictors (both continuous and categorical, linear and nonlinear) it still has important limitations. First, fitting separate models for each datapoint ignores the spatiotemporal dependencies inherent in M/EEG data, potentially reducing statistical power and interpretability. Second, the subsequent group-level analyses typically do not account for hierarchical dependencies which could otherwise be addressed through multilevel modelling. Finally, because the output of this procedure is summarised by cluster-based inference, its conclusions remain subject to the limitations discussed in the previous section.

Beyond modelling nonlinear effects of continuous predictors on ERP amplitudes, GAMs have been employed to capture the temporal dynamics of ERPs themselves, effectively modelling the shape of the waveform over time (Abugaber et al., 2023; Baayen et al., 2018; Meulman et al., 2015, 2023). This approach allows for the estimation of smooth, data-driven functions that characterise how neural responses evolve over time, offering a flexible alternative to traditional linear models. These developments show that regression models can move M/EEG analysis beyond simple averaging and sample-by-sample testing. GAMs provide a particularly natural extension because they model the temporal trajectory itself as a smooth, data-driven function.

### 1.5 Generalised additive models

In generalised additive models, the functional relationship between the predictors and the response variable is decomposed into a sum of low-dimensional non-parametric functions. A typical GAM has the following form:

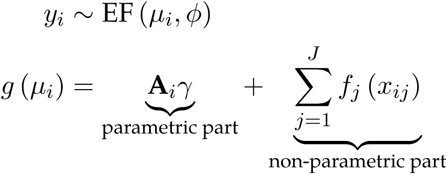

where *y_i_ ∼* EF (*µ_i_, ϕ*) denotes that the observations *y_i_* are distributed as some member of the exponential family of distributions (e.g., Gaussian, Gamma, Beta, Poisson) with mean *µ_i_* and scale parameter *ϕ*; *g*(*·*) is the link function, **A***_i_* is the *i*th row of a known parametric model matrix, *γ* is a vector of parameters for the parametric terms (to be estimated), *f_j_* is a smooth function of covariate *x_j_* (to be estimated as well). The smooth functions *f_j_* are represented in the model as a weighted sum of *K* simpler, basis functions:

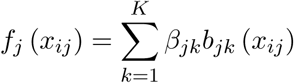

where *β_jk_* is the weight (coefficient) associated with the *k*th basis function *b_jk_*() evaluated at the covariate value *x_ij_* for the *j*th smooth function *f_j_*. To clarify the terminology at this point: *splines* are functions composed of simpler functions. These simpler functions are called *basis functions* (e.g., cubic polynomial, thin-plate) and the set of basis functions is called a *basis*. Each basis function is weighted by its coefficient and the resultant spline is the sum of these weighted basis functions (Figure 1A). Spline coefficients are penalised (usually through the square of the smooth functions’ second derivative) in a way that can be interpreted, in Bayesian terms, as a prior on the “wiggliness” of the function (Miller, 2025; Wood, 2017). In other words, more complex (wiggly) basis functions are automatically penalised.

**Figure 1.**
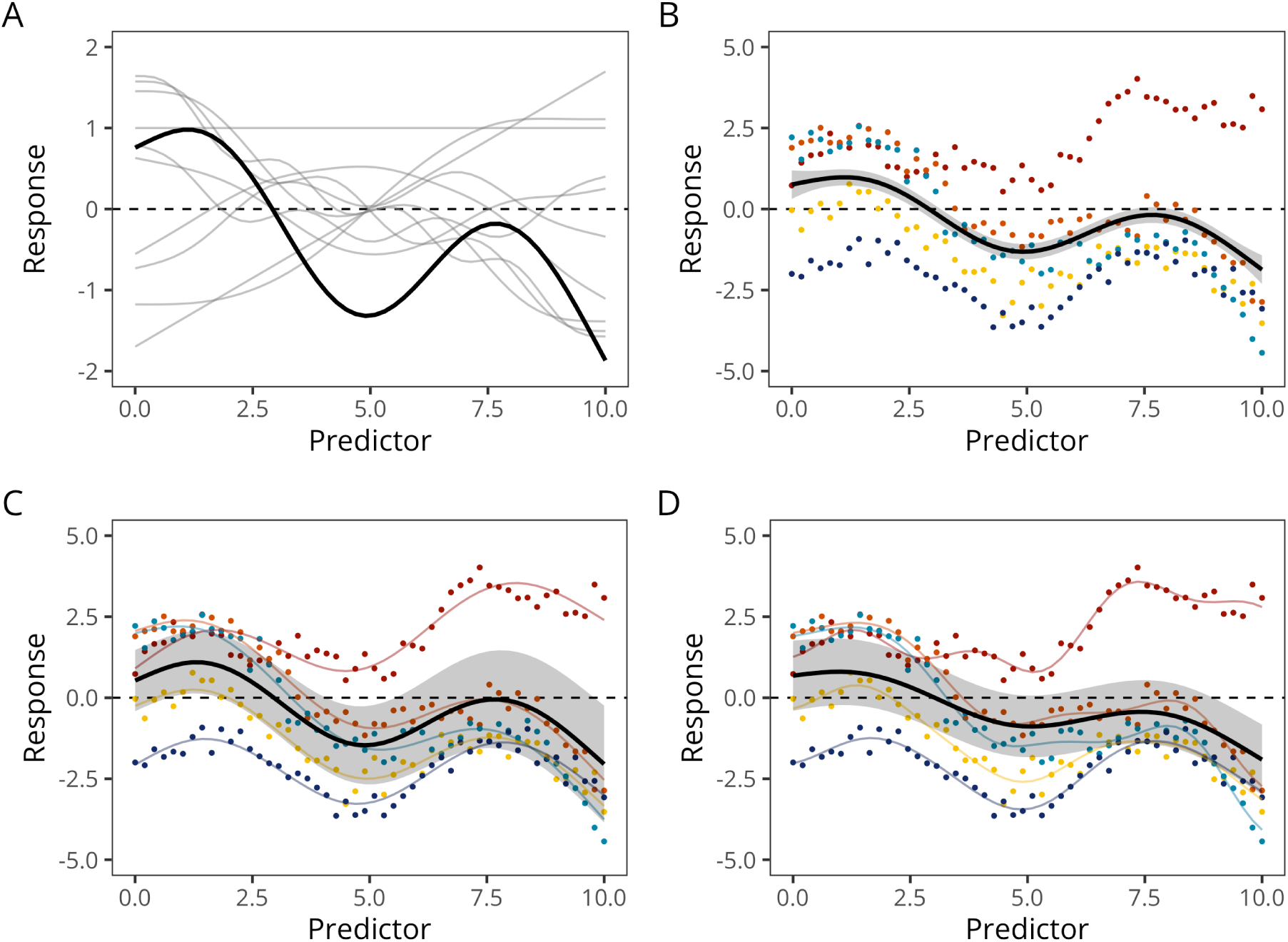
Different types of GAMs. **A**: GAMs predictions are computed as the weighted sum (in black) of basis functions (here thin-plate basis functions, in grey). **B**: Constant-effect GAM, with 5 participants in colours and the group-level prediction in black. **C**: Varying-intercept + varying-slope GAMM (with constant smoother). **D**: Varying-intercept + varying-slope + varying-smoother GAMM. In this model, each participant gets its own intercept, slope, and degree of ‘wiggliness’ (smoother).

A detailed treatment of the technical underpinnings of GAMs is beyond the scope of this article (see reference books such as Hastie & Tibshirani, 2017; Wood, 2017). However, it is worth emphasising that GAMs have been successfully applied to a wide range of time series data across the cognitive sciences, including pupillometry (e.g., Rij et al., 2019), articulography (e.g., Wieling, 2018), speech formant dynamics (e.g., Sóskuthy, 2021), neuroimaging data (e.g., Dinga et al., 2021), and event-related potentials (e.g., Abugaber et al., 2023; Baayen et al., 2018; Meulman et al., 2015, 2023). Their appeal for modelling M/EEG data lies in their ability to flexibly capture the complex shape of ERP waveforms without overfitting, through the use of smooth functions constrained by penalisation. Recent extensions, such as distributional GAMs (Rigby & Stasinopoulos, 2005; Umlauf et al., 2018), allow researchers to model not only the mean structure but also the variance (or scale) and other distributional properties as functions of predictors, a feature that has proven useful in modelling neuroimaging data (e.g., Dinga et al., 2021). Moreover, hierarchical or multilevel GAMs (E. J. Pedersen et al., 2019) provide a principled way to account for the nested structure of M/EEG data (e.g., trials within participants), enabling the inclusion of varying intercepts, slopes, and smoothers (as illustrated in Figure 1C-D). This approach mitigates the risk of overfitting and reduces the influence of outliers on smooth estimates (Baayen & Linke, 2020; Meulman et al., 2023).

### 1.6 Objectives

Here, we develop and evaluate a model-based framework for temporally localised inference in M/EEG data using Bayesian generalised additive multilevel models implemented in brms (Bürkner, 2017, 2018). We first benchmark the complete analysis workflow against commonly used point-wise, multiplicity-corrected, cluster-based, and change point procedures using simulated EEG data with known temporal ground truth across four SNR levels. We then apply the same framework to empirical MEG decoding data and evaluate the stability of temporal detections using repeated split-half resampling. Our goal is not to isolate the independent contribution of smoothing, Bayesian inference, or multilevel modelling, but to assess the operating characteristics of the complete model-based workflow.

## 2 Benchmarking with known ground truth

### 2.1 Methods

#### 2.1.1 M/EEG data simulation

To assess the accuracy of group-level temporal localisation with our proposed method, we simulated EEG data with known ground-truth temporal effects. Following the general approach of Sassenhagen & Draschkow (2019) and Rousselet (2025), each simulated dataset comprised two experimental conditions: a reference condition containing noise only and a target condition containing the same background activity plus a time-localised signal. Each dataset contained 20 participants, with 50 trials per participant and condition, sampled at 500 Hz over a 0.5-s epoch. The simulation framework allowed the temporal structure of the true effect to be specified explicitly. The true effect extended from 160 to 340 ms.

To introduce participant-level heterogeneity, the amplitude of the true effect was scaled separately for each participant, with scaling coefficients drawn from a truncated normal distribution with mean 1 and standard deviation 0.25. Participant-level noise magnitude was also varied, using a truncated normal distribution with mean 1 and standard deviation 0.5. Signal strength was manipulated across four signal-to-noise ratio (SNR) levels (0.25, 0.5, 1, 2). SNR scaling was referenced to the 0-100 ms baseline period, which was also used subsequently to derive the baseline-dependent thresholds for the BGAM analyses. Figure 2 illustrates one simulated dataset.

**Figure 2.**
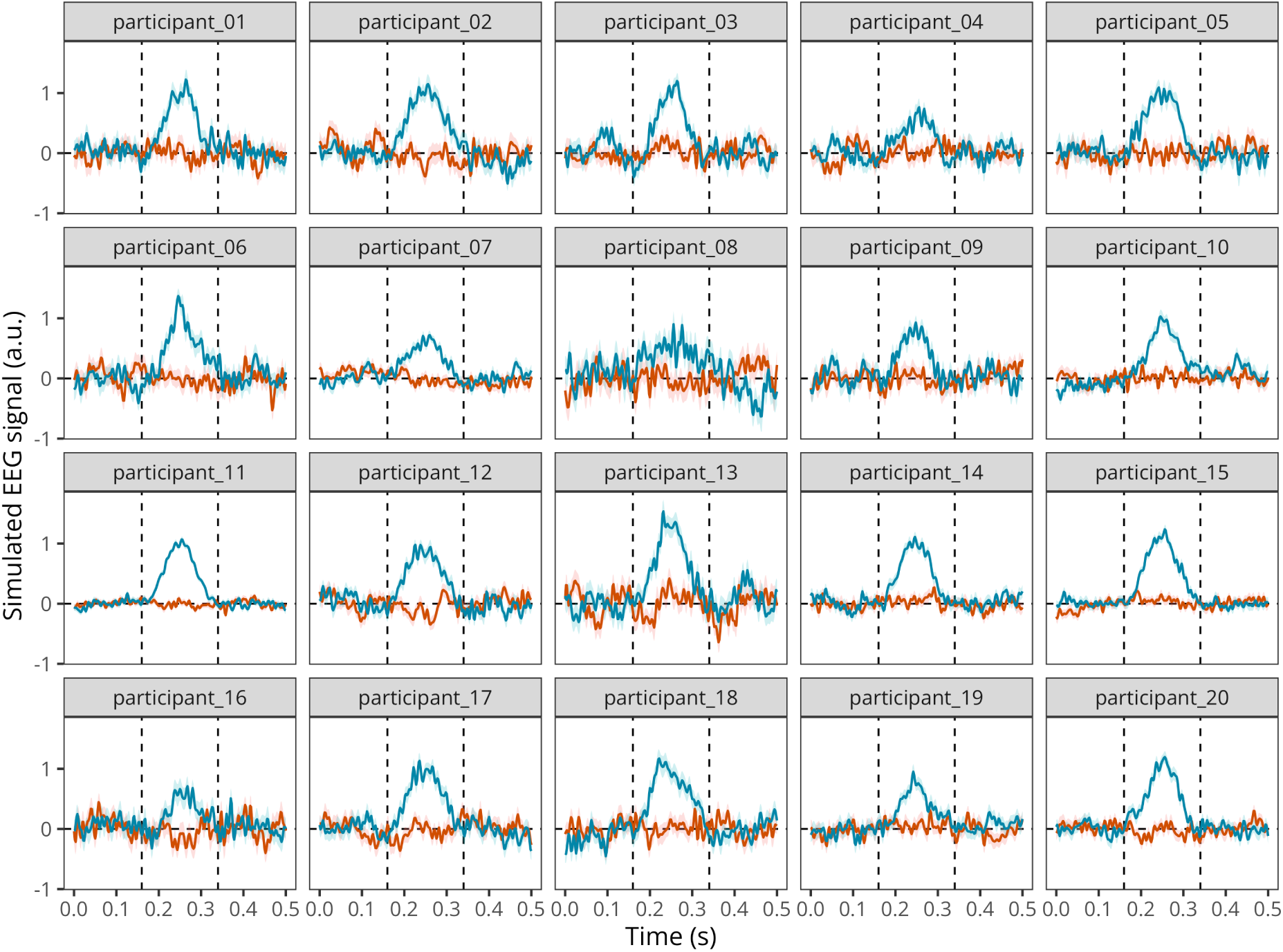
Mean simulated EEG activity in two conditions with 50 trials each, for a group of 20 participants. The error band represents the mean +/− 1 standard error of the mean.

#### 2.1.2 Model description and model fltting

We first fitted a Bayesian generalised additive model (BGAM) using the brms package (Bürkner, 2017, 2018), which we subsequently extended to a Bayesian generalised additive multilevel model (BGAMM) to account for between-participant variability (for an introduction to Bayesian multilevel modelling in brms, see Nalborczyk et al., 2019). The BGAMM included population-level (i.e., “fixed”) intercepts, slopes, and smooth effects of time, as well as participant-level (i.e., “random”) varying intercepts, slopes, and smooths. Although this model can be fitted directly to single-trial data, we adopted a computationally more efficient formulation based on participant-level summary statistics (means and standard errors), analogous to a Bayesian meta-analytic model. Specifically, the reported standard errors were incorporated into the likelihood to account for differences in measurement precision across observations, while an additional residual variance parameter was estimated to capture variability not explained by these standard errors or the model predictors.

Posterior distributions were approximated using Markov chain Monte Carlo (MCMC) sampling, with four independent chains of 5,000 iterations each, including 1,000 warmup iterations per chain. This yielded a total of 16,000 post-warmup draws for posterior inference. Convergence was assessed through visual inspection of trace plots and the rank-normalised split-*R*^ diagnostic (Gabry et al., 2019; Gelman et al., 2020; Vehtari et al., 2021a).

Figure 3 illustrates the BGAMM predictions alongside the simulated data. The model adequately captures the underlying temporal dynamics of both conditions (see additional posterior predictive checks in Appendix C). Panel A shows the posterior expected responses for each condition, while Panel B shows the estimated time-varying condition contrast *β*(*t*). The shaded regions represent point-wise 95% credible intervals, which quantify posterior uncertainty at each time point, conditional on the model, priors, and observed data.

**Figure 3.**
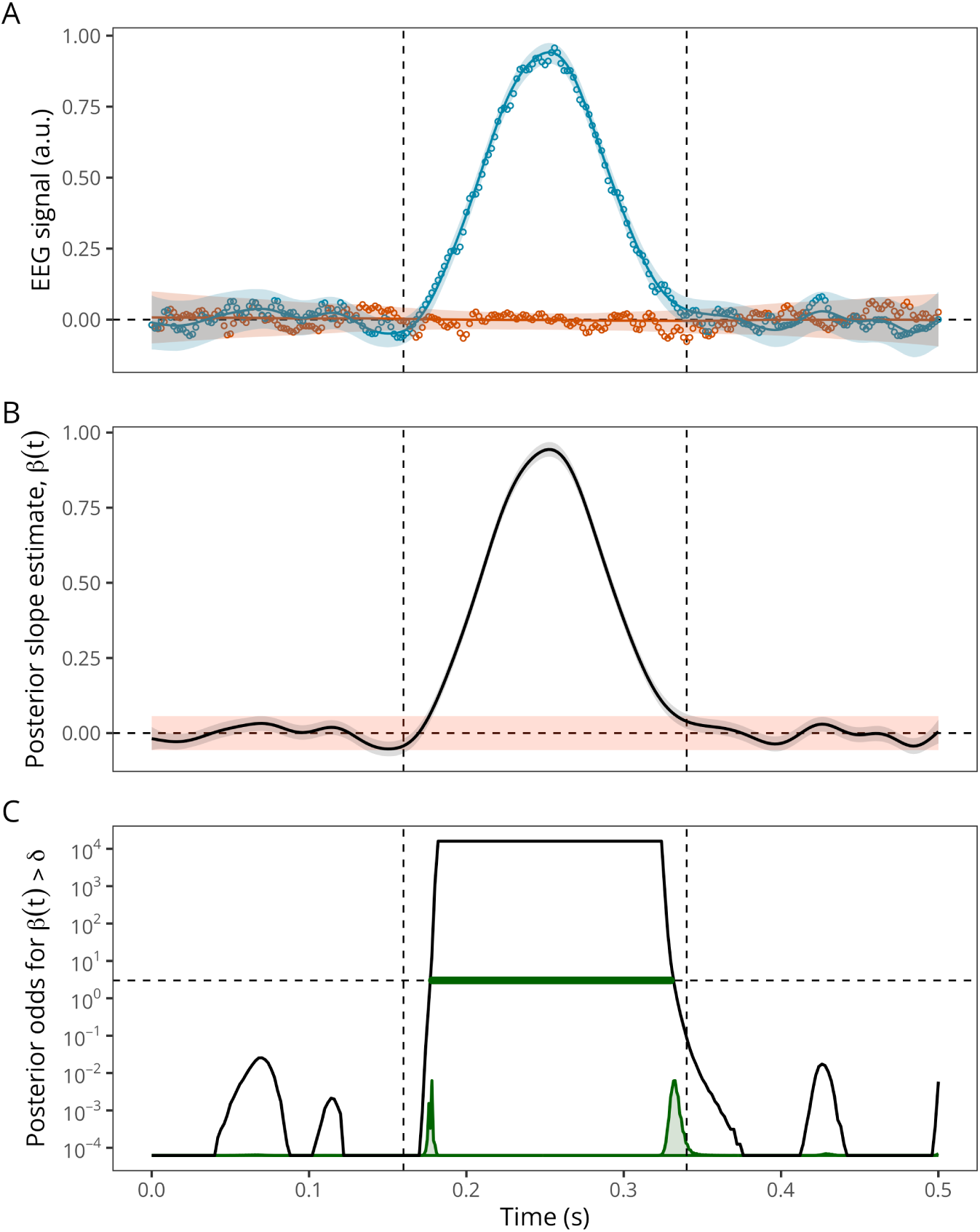
Illustration of BGAMM-based estimation and inference on simulated EEG data. (A) Observed signals (points) and posterior predictions (solid lines), with shaded areas representing point-wise 95% credible intervals for each condition. (B) Posterior median of the time-varying condition contrast with its point-wise 95% credible interval (gray shading). The orange region represents the region of practical equivalence (ROPE), defined symmetrically around zero using a baseline-derived smallest effect size of interest (SESOI), corresponding to the maximum absolute bound of the point-wise 95% credible interval of the condition contrast during the baseline interval (0-0.1 s). (C) Posterior odds that the posterior contrasts exceeds the SESOI, displayed on a logarithmic scale. The horizontal dashed line indicates the decision threshold (posterior odds = 3), with periods exceeding this threshold highlighted in green. The distributions near the bottom represent the posterior uncertainty in effect onset and offset, estimated independently from threshold crossings in individual posterior draws and rescaled for visualisation. Vertical dashed lines indicate the true simulated onset and offset of the effect. Posterior odds are bounded by the finite number of posterior draws.

To identify periods of substantial condition differences, we computed the posterior probability that the time-varying condition contrast, *β*(*t*), exceeds a smallest effect size of interest (SESOI), denoted *δ*. We operationally defined this SESOI from the pre-effect baseline interval (0-0.1 s) as the maximum absolute bound of the point-wise 95% credible intervals for the condition contrast, considering both their lower (2.5th percentile) and upper (97.5th percentile) bounds across all baseline time points. This definition ensures that the resulting symmetric region of practical equivalence [ROPE; Kruschke & Liddell (2017)], [*−δ, δ*], encompasses the entire point-wise 95% credible interval at every baseline time point (orange shading in Panel B). This baseline-derived threshold provides a model-based reference against which post-onset effects can be evaluated, accounting for both the magnitude and posterior uncertainty of estimated condition differences during a period when no effect is expected.

At each time point, we computed the posterior probability of exceeding this threshold, *p*(*t*) = Pr(*β*(*t*) *> δ |* data), and converted it into posterior odds, *p*(*t*)/(1 *− p*(*t*)). These odds quantify the relative posterior support for the condition contrast exceeding versus not exceeding the SESOI. For example, posterior odds of 3 indicate that an effect exceeding *δ* is three times as probable as one not exceeding it, conditional on the model and observed data. As illustrated in Panel C, contiguous time points with posterior odds exceeding a predefined threshold (3 in this example) were grouped into temporal clusters, providing estimates of the onset and offset of periods with strong posterior evidence for an effect. The operating characteristics of this procedure were evaluated in the simulation study. Because inference is based on posterior contrasts, the approach can be extended to a broad range of model structures and experimental designs.

In addition to identifying effect intervals from the posterior-odds trajectory, we quantified uncertainty in their temporal boundaries directly. For each joint posterior draw of the time-varying condition contrast, we identified the earliest and latest time points at which *β*(*t*) exceeded the same SESOI, *δ*. Repeating this procedure across posterior draws yielded distributions of effect onset and offset (illustrated near the bottom of Panel C). This provides complementary information to the posterior-odds-based approach: whereas the latter identifies periods with sufficiently strong posterior evidence for an effect, the former characterises posterior uncertainty about the timing of its boundaries.

#### 2.1.3 Comparing the onset/offset estimates across approaches

We then compared the ability of the BGAM to accurately estimate the onset and offset of the ERP difference to other widely-used methods. First, we conducted mass univariate t-tests (thus treating each time point independently) and identified the onset and offset of the ERP difference as the first and last values crossing an arbitrary significance threshold (*α* = 0.05). We then followed the same approach but after applying different forms of multiplicity correction to the *p*-values. We compared two methods that control the FDR (i.e., BH95, Benjamini & Hochberg, 1995; and BY01, Benjamini & Yekutieli, 2001), one method that controls the FWER (i.e., Holm– Bonferroni method, Holm, 1979), and two cluster-based permutation methods (permutation with a single cluster-forming threshold and threshold-free cluster enhancement, TFCE, S. Smith & Nichols, 2009). The BH95, BY01, and Holm corrections were applied to the *p*-values using the p.adjust() function in R. The cluster-based inference was implemented using a cluster-sum statistic of *t*-values, as implemented in MNE-Python (Gramfort, 2013), called via the R package reticulate v 1.46.0 (Ushey et al., 2024).

We also compared these estimates to the onset and offset estimated using the binary segmentation algorithm, as implemented in the R package changepoint v 2.3 (Killick et al., 2022), and applied directly to the squared *t*-values (Rousselet, 2025). In contrast to the participant-level application evaluated by Rousselet (2025), we applied this procedure to the group-level statistic, and allowed the number of change points to be selected automatically. We therefore treat it as a related change point comparator rather than a direct replication of Rousselet (2025)’s implementation.

Importantly, these comparisons evaluate the complete proposed workflow, which combines temporal smoothing, multilevel estimation, and Bayesian posterior inference, against commonly used implementations of the alternative procedures. They are therefore not intended to isolate the independent contribution of smoothing, Bayesian inference, or multilevel modelling to any observed performance difference.

Figure 4 illustrates the temporal detections obtained by the alternative procedures for one simulated dataset. The methods differ both in the extent and fragmentation of their detections: several corrected procedures identify intervals largely contained within the true effect, whereas the uncorrected and FDR-based point-wise procedures also yield detections outside its temporal boundaries. This example is intended to illustrate the behaviour of the different procedures; their operating characteristics across simulations are quantified below.

**Figure 4.**
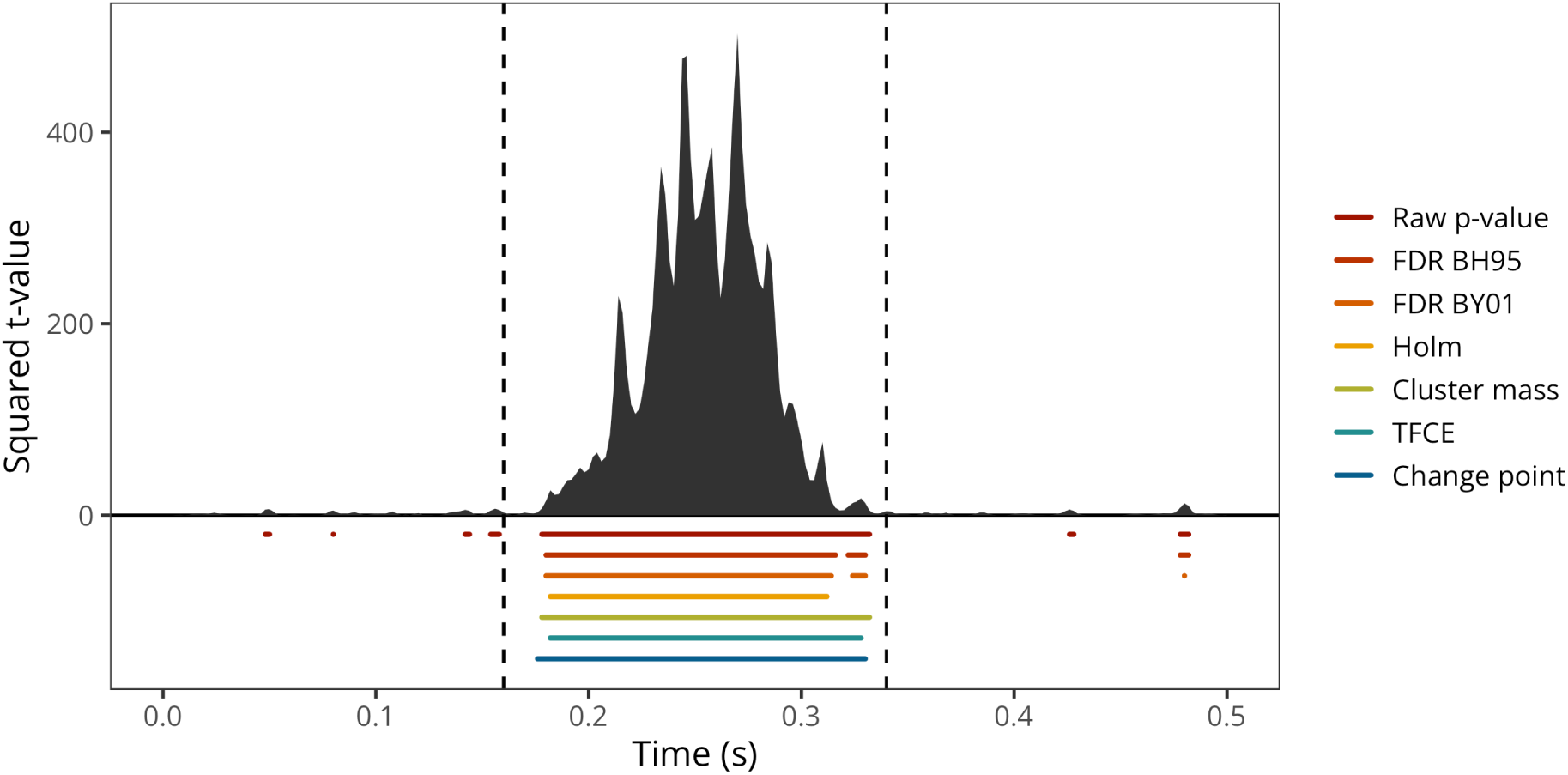
Example time course of squared t-values in simulated data. The grey area shows the squared t-statistic over time, and the vertical dashed lines mark the true simulated onset and offset of the effect. Coloured horizontal segments indicate temporal clusters detected by each method: uncorrected p-values, corrected p-values (FDR BH95, FDR BY01, Holm), cluster-based inference (cluster mass, TFCE), and binary segmentation change point detection.

#### 2.1.4 Simulation study

To assess the accuracy of group-level temporal localisation, all methods were evaluated across 250 simulated datasets at each of four SNR levels, yielding 1,000 simulated datasets in total.

For each simulated dataset and method, the complete set of detected time intervals was compared with the true effect interval(s). Overall agreement was quantified using temporal intersection-over-union (IoU; equivalently, the Jaccard index), defined as the duration of the intersection between the detected and true interval sets divided by the duration of their union. IoU therefore jointly penalises both missed portions of the true effect and detections extending outside the true effect. Performance was summarised across simulations using the median IoU, with values closer to 1 indicating more accurate recovery of the true temporal extent.

Boundary localisation was evaluated separately for each true effect interval. Within the entire window, all detected intervals were considered jointly: the estimated onset was defined as the earliest detected onset and the estimated offset as the latest detected offset. This deliberately stringent definition penalises fragmented or spurious detections rather than selecting only the detected interval that best matches the true effect. Onset and offset errors were calculated as the estimated minus true boundary, such that negative values indicate boundaries estimated too early and positive values indicate boundaries estimated too late. For recovered true clusters, boundary localisation was summarised across simulations using the mean signed error (bias), mean absolute error (MAE), root mean squared error (RMSE), and standard deviation (SD) of the signed errors, all expressed in milliseconds.

These simulations were designed to evaluate repeated-sampling temporal-recovery performance under fixed, controlled data-generating scenarios. They were not designed as a simulation-based calibration study and the results should therefore not be interpreted as evidence of prior or posterior calibration.

### 2.2 Results

Across SNRs, BGAM consistently achieved the highest median temporal overlap with the true effect, with IoU increasing from 0.75 at SNR = 0.25 to 0.92 at SNR = 2 (Figure 5A). Boundary estimates were also comparatively well centred: BGAM showed a small positive onset bias and offset bias close to zero across SNRs (Figure 5B). The change point method yielded similarly small signed boundary biases, but lower temporal overlap, indicating that accurate average boundary location did not necessarily translate into accurate recovery of the full effect interval. In contrast, raw point-wise testing markedly overextended the detected effect, with systematically early onsets and late offsets, whereas several corrected frequentist procedures (particularly Holm, TFCE, FDR BY01, and cluster mass) tended to show the opposite pattern at lower SNRs, with delayed onsets and premature offsets consistent with temporal contraction. As SNR increased, differences in IoU between methods narrowed, although BGAM retained the highest overlap with the true temporal extent.

**Figure 5.**
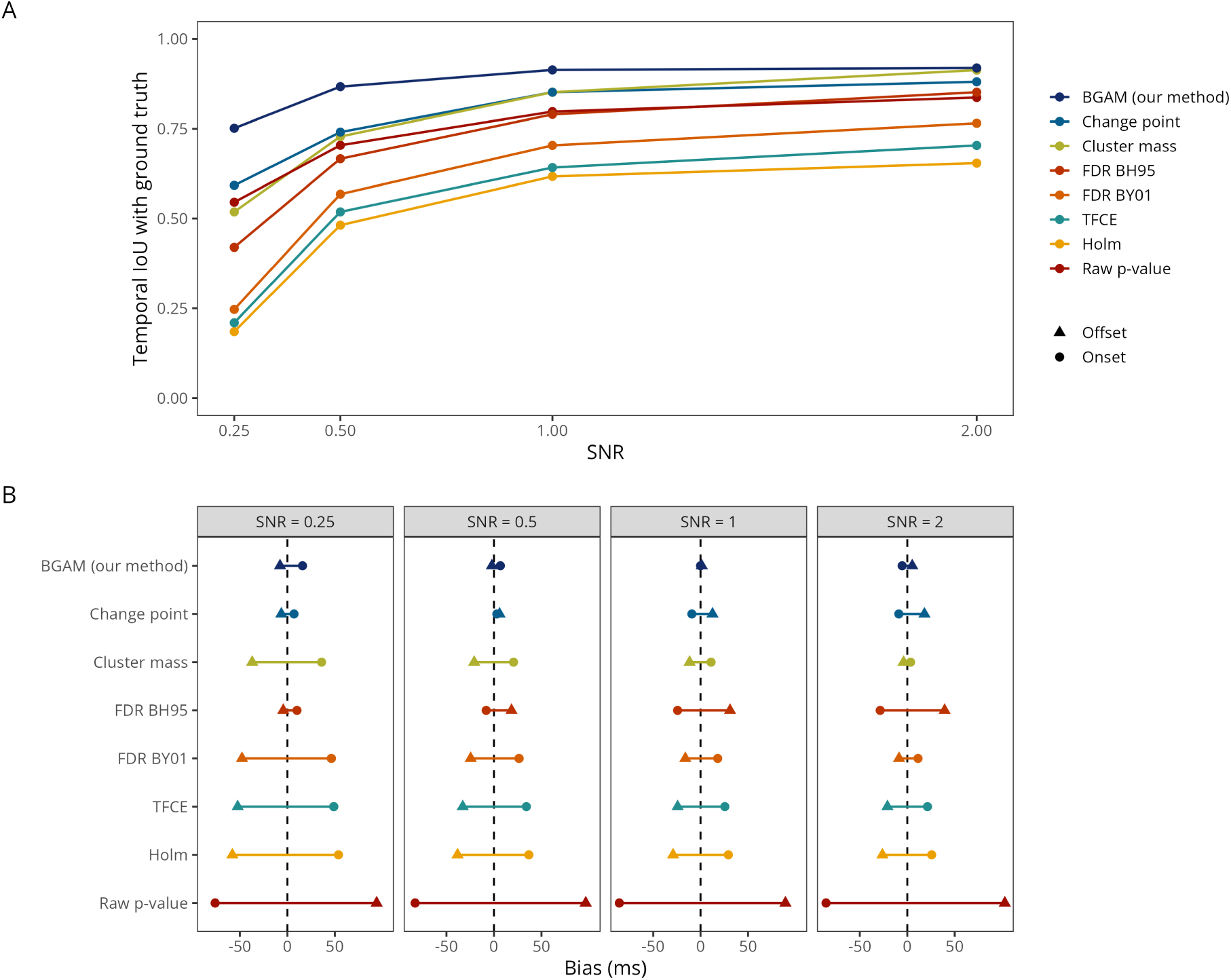
Temporal detection and boundary localisation performance. (A) Median temporal intersection-over-union (IoU) between the detected and true effect intervals across simulations, as a function of signal-to-noise ratio (SNR). Higher IoU indicates greater agreement between the detected and true temporal extent of the effect. (B) Mean signed localisation bias of the estimated onset (circles) and offset (triangles) boundaries. Negative values indicate that a boundary was estimated earlier than its true location, whereas positive values indicate that it was estimated later. In panel B, methods are ordered by their average absolute onset/offset bias across SNRs, with lower-error methods shown toward the top.

The results are further summarised in Table 1. Across all SNR levels, BGAM achieved the highest median temporal overlap with the true effect, with intersection-over-union (IoU) increasing from 0.75 at SNR = 0.25 to 0.92 at SNR = 2. This advantage was particularly pronounced at lower SNRs, whereas differences between methods narrowed as signal strength increased. BGAM also provided comparatively accurate onset localisation, with mean absolute error (MAE) decreasing from 20.1 ms at SNR = 0.25 to 7.9 ms at SNR = 2. Its onset estimates showed relatively small systematic bias, approaching zero at intermediate SNRs. At the highest SNR, cluster-mass inference achieved marginally lower onset MAE and RMSE, although the differences were small.

**Table 1.** Temporal detection and boundary localisation performance across signal-to-noise ratios (SNRs). For each method and SNR, onset and offset localisation are summarised using mean signed bias, mean absolute error (MAE), root mean squared error (RMSE), and standard deviation (SD), all expressed in milliseconds. Negative bias indicates a boundary estimated earlier than its true location, whereas positive bias indicates a later estimate. Temporal overlap is quantified using the median intersection-over-union (IoU) between the detected and true effect intervals across simulations, with higher values indicating better recovery of the true temporal extent. Bold values indicate the best-performing method for each metric within each SNR; for signed bias, performance is evaluated using absolute bias.

|  | Onset |  |  |  | Offset |  |  |  | Temporal overlap |
|---|---|---|---|---|---|---|---|---|---|
| Method | Bias | MAE | RMSE | SD | Bias | MAE | RMSE | SD | IoU |
| SNR = 0.25 |  |  |  |  |  |  |  |  |  |
| BGAM (our method) | 16.02 | **20.06** | **23.43** | **17.14** | -7.76 | **29.63** | 46.62 | 46.06 | **0.75** |
| Change point | **7.02** | 42.91 | 56.11 | 55.78 | -6.40 | 46.00 | 60.37 | 60.15 | 0.59 |
| Raw p-value | -76.14 | 85.13 | 99.40 | 64.03 | 94.14 | 100.30 | 116.23 | 68.30 | 0.55 |
| Cluster mass | 36.15 | 38.71 | 41.66 | 20.74 | -36.98 | 40.00 | **43.32** | 22.61 | 0.52 |
| FDR BH95 | 10.14 | 48.99 | 59.16 | 58.40 | **-4.35** | 56.62 | 68.84 | 68.84 | 0.42 |
| FDR BY01 | 46.39 | 52.33 | 55.13 | 29.85 | -47.75 | 54.28 | 57.43 | 31.98 | 0.25 |
| TFCE | 48.91 | 53.42 | 55.87 | 27.07 | -52.29 | 56.34 | 58.89 | 27.14 | 0.22 |
| Holm | 53.80 | 56.31 | 58.25 | 22.38 | -57.93 | 59.07 | 60.99 | **19.14** | 0.19 |
| SNR = 0.5 |  |  |  |  |  |  |  |  |  |
| BGAM (our method) | 6.78 | **11.16** | **13.61** | **11.83** | **-2.18** | **14.88** | 27.65 | 27.62 | **0.86** |
| Change point | **3.06** | 29.06 | 41.88 | 41.85 | 5.91 | 35.91 | 55.75 | 55.55 | 0.75 |
| Cluster mass | 20.68 | 22.07 | 24.53 | 13.22 | -20.87 | 21.75 | **23.96** | **11.78** | 0.73 |
| Raw p-value | -83.10 | 88.18 | 101.73 | 58.80 | 96.59 | 102.22 | 117.29 | 66.67 | 0.71 |
| FDR BH95 | -8.22 | 39.21 | 53.52 | 52.99 | 18.54 | 47.90 | 66.60 | 64.10 | 0.67 |
| FDR BY01 | 26.42 | 31.78 | 35.08 | 23.13 | -24.60 | 34.79 | 41.22 | 33.15 | 0.57 |
| TFCE | 34.05 | 36.00 | 38.01 | 16.94 | -32.95 | 36.84 | 39.94 | 22.61 | 0.51 |
| Holm | 36.70 | 38.62 | 40.48 | 17.11 | -38.34 | 39.25 | 40.85 | 14.14 | 0.48 |
| SNR = 1 |  |  |  |  |  |  |  |  |  |
| BGAM (our method) | **-0.01** | **8.44** | **11.01** | 11.03 | **1.25** | **9.41** | 19.89 | 19.89 | **0.91** |
| Cluster mass | 10.95 | 12.06 | 14.16 | **8.99** | -11.54 | 12.54 | **14.49** | **8.78** | 0.85 |
| Change point | -9.19 | 26.47 | 48.07 | 47.28 | 12.47 | 29.70 | 55.34 | 54.02 | 0.83 |
| Raw p-value | -85.61 | 88.42 | 102.85 | 57.11 | 89.28 | 93.57 | 111.44 | 66.82 | 0.79 |
| FDR BH95 | -24.25 | 41.99 | 62.61 | 57.84 | 30.97 | 48.95 | 72.32 | 65.48 | 0.78 |
| FDR BY01 | 17.95 | 24.88 | 29.62 | 23.61 | -16.08 | 25.74 | 32.64 | 28.46 | 0.70 |
| TFCE | 25.46 | 28.65 | 31.63 | 18.79 | -24.19 | 28.86 | 32.61 | 21.91 | 0.64 |
| Holm | 29.18 | 31.19 | 33.58 | 16.67 | -28.99 | 29.95 | 31.58 | 12.56 | 0.61 |
| SNR = 2 |  |  |  |  |  |  |  |  |  |
| BGAM (our method) | -5.44 | 7.90 | 10.22 | **8.67** | 5.21 | 7.59 | 12.24 | 11.09 | **0.92** |
| Cluster mass | **3.50** | **7.70** | **9.60** | 8.96 | **-3.93** | **7.37** | **9.11** | **8.24** | 0.91 |
| Change point | -8.97 | 19.18 | 38.41 | 37.43 | 18.11 | 26.42 | 51.92 | 48.76 | 0.87 |
| FDR BH95 | -28.48 | 38.83 | 60.28 | 53.23 | 39.42 | 48.67 | 75.93 | 65.02 | 0.85 |
| Raw p-value | -85.46 | 86.63 | 102.11 | 55.99 | 102.82 | 103.71 | 119.20 | 60.43 | 0.83 |
| FDR BY01 | 11.30 | 21.18 | 28.58 | 26.31 | -8.72 | 23.10 | 33.21 | 32.11 | 0.77 |
| TFCE | 21.32 | 23.59 | 25.76 | 14.49 | -20.82 | 23.66 | 26.11 | 15.79 | 0.71 |
| Holm | 25.73 | 26.77 | 28.36 | 11.96 | -26.12 | 26.25 | 27.50 | 8.61 | 0.66 |

Offset localisation showed a more nuanced pattern. BGAM yielded the lowest offset MAE at SNRs of 0.25, 0.5, and 1, with offset bias remaining relatively close to zero across all SNRs. However, offset estimates were generally more variable than onset estimates, and cluster-mass inference consistently achieved lower offset RMSE, suggesting that BGAM was more susceptible to occasional large offset errors. In contrast, uncorrected point-wise testing systematically overes-timated the temporal extent of the effect, producing substantially earlier onsets and later offsets. Several multiplicity-corrected procedures, particularly Holm and TFCE, showed the opposite pattern, with delayed onsets and premature offsets, especially at lower SNRs. The change point method produced relatively small average boundary biases but substantially greater absolute errors and lower temporal overlap, illustrating that low signed bias does not necessarily imply accurate or reliable temporal localisation. Overall, these simulations indicate that BGAM provides particularly accurate recovery of the temporal extent and onset of the simulated effects, while its advantages are less consistent for offset localisation.

## 3 Application to empirical MEG data

### 3.1 Methods

To complement the ground-truth simulations, we applied the methods to an empirical MEG dataset from 32 participants performing a reading task (Nalborczyk et al., in preparation). Time-resolved multivariate pattern analysis yielded, for each participant, a decoding-accuracy time course expressed as ROC AUC. To test whether the group-level average decoding accuracy was above chance (i.e., 0.5) at each time point, we fitted a BGAMM as introduced previously with a basis dimension *k* = 30 and retained all time points exceeding a posterior odds of 10. To better distinguish signal from noise, we defined a baseline-derived threshold as the maximum absolute bound for the above-chance effect (AUC - 0.5) during the pre-stimulus baseline period. Although we chose a basis dimension of *k* = 30, which seemed appropriate for the present data, this choice should be adapted according to the properties of the modelled data (e.g., signal-to-noise ratio, prior low-pass filtering, sampling rate) and should be assessed by the usual model checking tools (e.g., model comparison, posterior predictive checking, see Appendix C).

To assess the stability of the temporal detections, we performed 1,000 random split-half repetitions of the 32 participants. For each repetition, participants were randomly divided into two non-overlapping groups of 16, and all methods were applied independently to each half. Temporal reproducibility was quantified using the intersection-over-union (IoU) between the sets of time points detected in the two halves, defined as the number of time points detected in both halves divided by the number detected in either half. Thus, an IoU of 1 indicates identical temporal detections, whereas an IoU of 0 indicates no temporal overlap. We additionally quantified agreement in temporal segmentation by recording whether the two halves yielded the same number of detected intervals. Because the repeated partitions reuse participants and are therefore not statistically independent, these resampling distributions were interpreted descriptively as measures of stability rather than as independent observations for inferential testing.

### 3.2 Results

Figure 6 shows the group-level average decoding performance through time together with the temporal intervals identified by each method. The Raw p-value and FDR BH95 procedures produced the most temporally extensive and fragmented detections, including above-chance intervals before stimulus onset and multiple late detections extending well beyond the main decoding response. The more conservative multiplicity-corrected and cluster-based procedures (FDR BY01, Holm, Cluster mass, and TFCE) identified a broadly similar main period of above-chance decoding beginning at approximately 50-60 ms, although some methods additionally yielded shorter, later clusters. In contrast, the Change point and BGAM procedures produced comparatively compact, contiguous intervals: the change-point interval extended from approximately +60 to +450 ms, while the BGAM detection covered a similar early period but ended earlier. Thus, the methods agreed broadly on the onset of the main decoding effect but differed substantially in its estimated duration and in the degree of temporal fragmentation.

**Figure 6.**
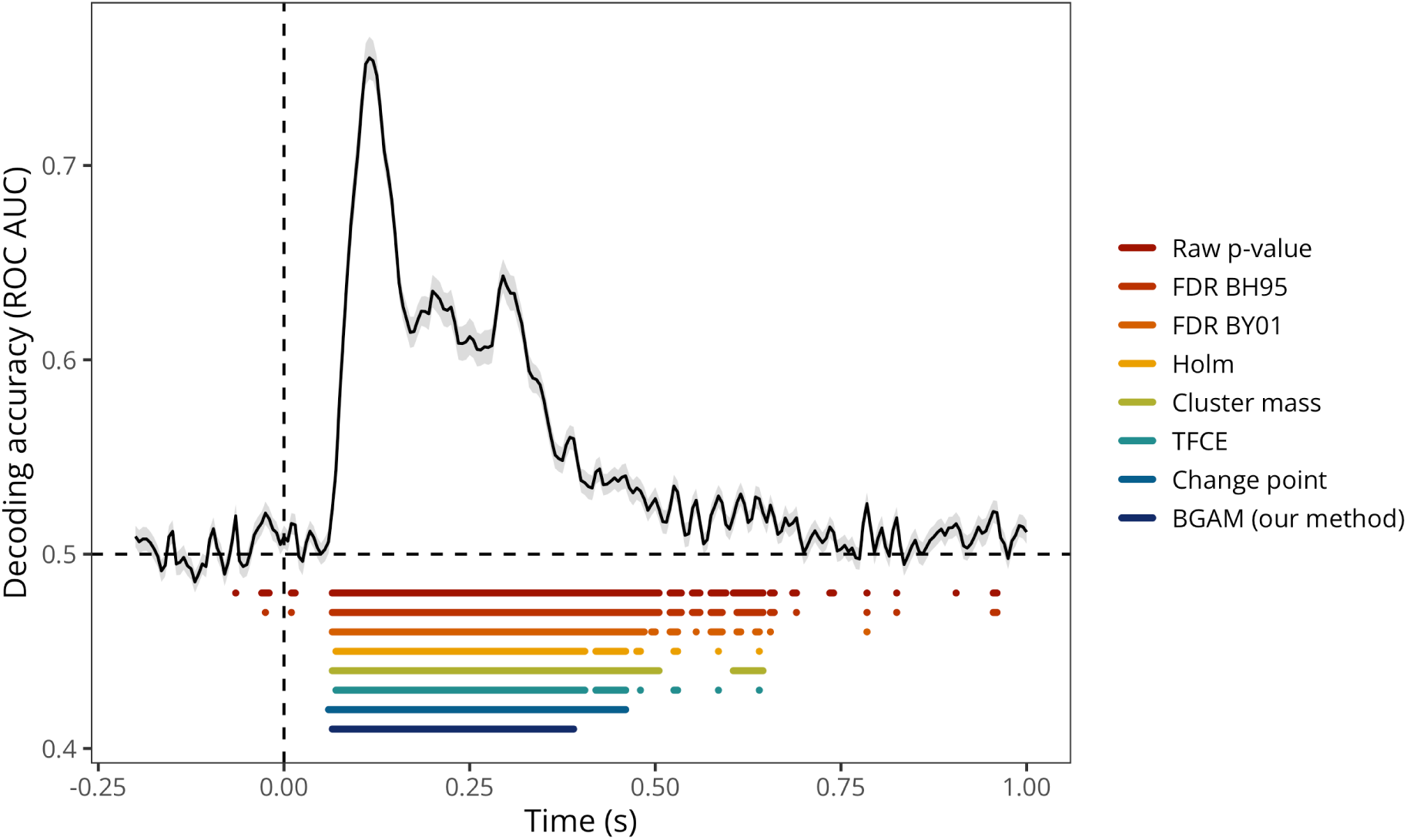
Group-level decoding performance and method-speciflc temporal detections. The black line shows mean decoding accuracy over time, expressed as ROC AUC, with the grey ribbon indicating the standard error of the mean. The horizontal dashed line marks chance-level performance, and the vertical dashed line marks stimulus onset. Coloured horizontal segments show time windows identified as above chance by each method. Data are from Nalborczyk et al. (in preparation).

Temporal detections were reproducible across random split halves, although stability varied substantially across methods. BGAM showed the greatest temporal overlap between halves (median IoU = 0.93), followed by cluster mass (0.87) and the change-point method (0.75); the remaining point-wise and multiplicity-corrected procedures yielded lower median overlap (0.60-0.72) (Figure 7A). BGAM also showed the most stable cluster count, with the two halves yielding the same number of detected intervals in almost all split-half repetitions (Figure 7B). Cluster-count agreement was substantially lower for the other procedures, including cluster-mass correction. When cluster counts differed between halves, discrepancies were approximately balanced between splits A and B, providing no indication of systematic split-specific asymmetry. Overall, BGAM combined the highest temporal overlap with highly consistent segmentation across repeated non-overlapping halves of the empirical sample.

**Figure 7.**
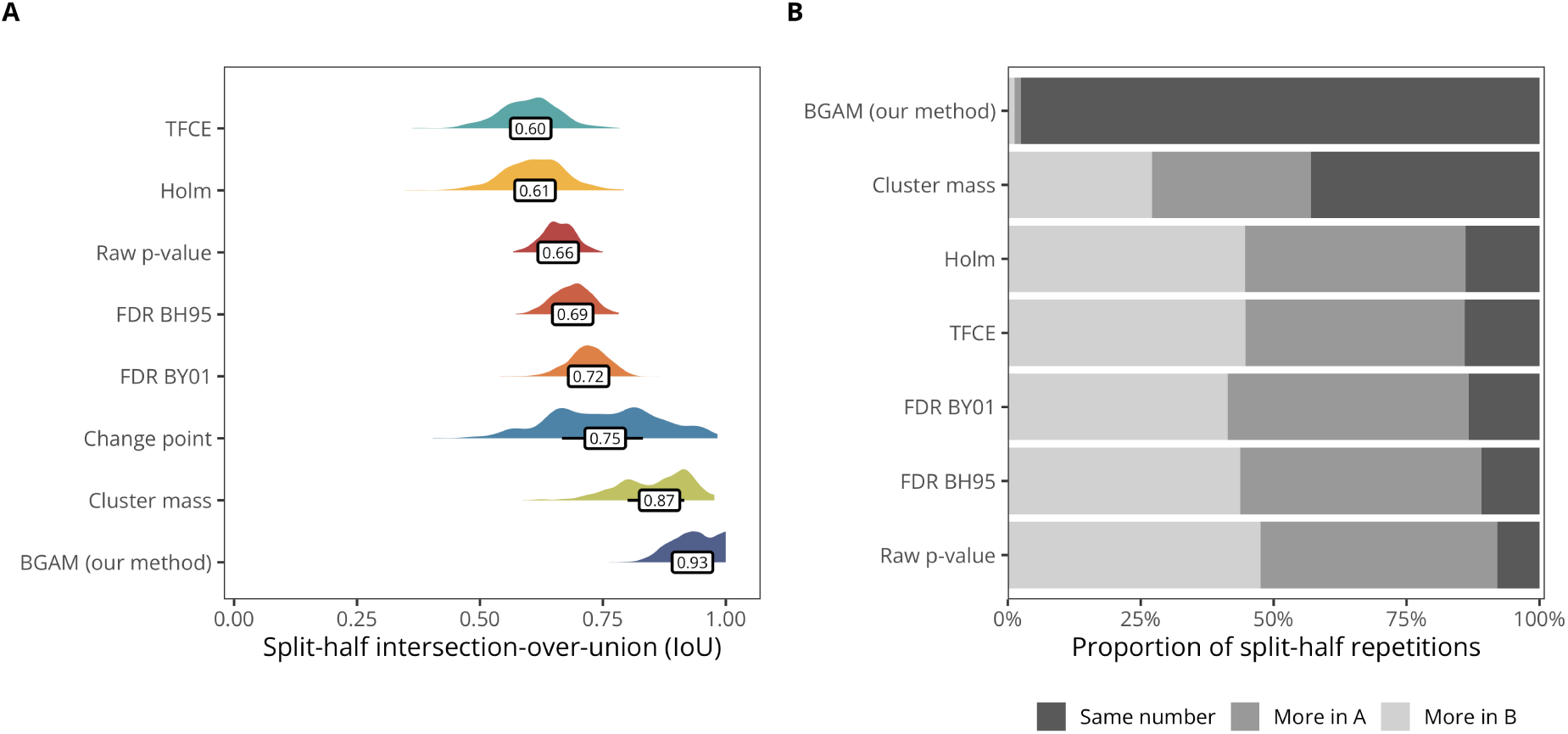
Split-half reliability of temporal detections in the empirical MEG data. (A) Distribution of temporal intersection-over-union (IoU) values across repeated random split-half analyses for each method. IoU quantifies the overlap between the sets of detected time intervals identified independently in the two halves, with higher values indicating greater temporal reproducibility. Labels indicate the median IoU across repetitions. (B) Proportion of split-half repetitions in which the two halves yielded the same number of detected temporal clusters, or in which more clusters were detected in split A or split B. Similar proportions of ‘more in A’ and ‘more in B’ indicate that differences in cluster number were not systematically associated with either half.

## 4 Discussion

In brief, our results show that the model-based approach provides strong temporal localisation performance across both ground-truth simulations and empirical resampling analyses. In simulations, BGAM consistently yielded the highest temporal overlap with the true effect across SNRs, together with comparatively accurate onset localisation and small systematic boundary biases. Its advantage was less uniform for offset localisation: offset estimates were more variable, and cluster-mass correction performed better on some offset-specific error metrics, particularly RMSE. The empirical split-half analysis provided a complementary assessment of stability when the ground truth was unknown. Here, BGAM yielded the highest temporal overlap between independently analysed halves of the sample (median IoU = 0.93), exceeding cluster mass (0.87) and the change-point procedure (0.75), while also producing highly consistent cluster counts across repetitions. Thus, the simulation and empirical analyses converge in showing strong recovery of temporal support with BGAM, while also highlighting that performance should be assessed along several complementary dimensions rather than reduced to a single onset or offset metric.

As with previous simulation studies (e.g., Rousselet et al., 2008; Sassenhagen & Draschkow, 2019), results inevitably depend on design choices, including the specific cluster-forming algorithm and threshold (for cluster-based methods), the signal-to-noise ratio, and the potential degradation of temporal resolution introduced by preprocessing steps such as low-pass filtering. Temporal filtering should not be assumed to affect all inferential procedures identically. Because different methods already impose different forms of temporal regularisation or aggregation, preprocessing filters may interact with the subsequent statistical analysis and alter temporal localisation differently across methods. These results therefore illustrate that relative performance depends not only on the inferential framework itself, but also on the interaction between the procedure, signal characteristics, and preprocessing choices. TFCE, for example, yielded lower temporal overlap than the traditional cluster-mass procedure in the present simulations, consistent with previous observations that different forms of cluster inference can have distinct localisation properties (e.g., Rousselet, 2025; S. Smith & Nichols, 2009). Interestingly, Holm and TFCE produced similar temporal detections in the empirical dataset despite relying on very different inferential mechanisms.

Beyond these comparison-specific considerations, the BGAM itself relies on modelling assumptions that require careful evaluation. A critical requirement for any model-based approach is that the model must adequately capture the relevant features of the data-generating process. Misspecified models are likely to produce biased or unreliable onset/offset estimates. This underscores the importance of thorough model diagnostics, including posterior predictive checks, fit assessments, and model comparison (Gelman et al., 2020). An important and related methodological consideration concerns the selection of model hyperparameters, such as the number of basis functions and the threshold for posterior odds. Although our simulations suggest that these parameters influence the precision and reliability of onset and offset estimates, optimal values may vary depending on the signal’s temporal dynamics and signal-to-noise characteristics. Future work could explore principled approaches to hyperparameter tuning, including cross-validation or fully Bayesian model selection using tools such as leave-one-out cross-validation (Gelman et al., 2020). We provide initial guidance in Appendix C and advocate for future development of adaptive heuristics to support flexible yet parsimonious model specification. It should also be noted that penalised smooths are not designed to reproduce true discontinuities exactly; abrupt changes may be temporally smeared, particularly when evidence is weak, and the resulting boundary error can depend on both regularisation and the decision rule used to define an effect.

Beyond specification of the mean trajectory, a second modelling consideration concerns residual temporal dependence. The smooth terms used here model temporal structure in the conditional mean, but do not imply that the remaining residuals are temporally independent. If substantial residual autocorrelation remains, posterior uncertainty may be miscalibrated even when the mean trajectory is well described. The simulated data used here contain temporal dependence that is not represented by an explicit residual-correlation process, yet temporal recovery remained accurate under the conditions considered. Nevertheless, we recommend inspecting residual dependence in empirical applications and considering an explicit autocorrelation structure when warranted.

Other limitations concern the scope of the present implementation and simulations. The current approach estimates temporal effects independently at each sensor (1D temporal data). Extending the current framework to incorporate additional temporal (see Appendix B) or spatial dimensions would improve both sensitivity and interpretability. Such extensions could draw on methods from spatial epidemiology and geostatistics using either GAMMs or approximate Gaussian process regression (e.g., Rasmussen & Williams, 2005; Riutort-Mayol et al., 2023), depending on computational feasibility. Moreover, the present simulations primarily evaluate group-level inference with a multilevel model and therefore should not be taken as evidence of equivalent localisation performance when the model is fitted to a single participant, particularly in low-SNR settings where no between-participant pooling is available. Hierarchical bootstrap procedures provide another complementary route to population-level uncertainty in temporal summaries and may be preferable when the inferential target is a resampled population distribution of onset or offset estimates rather than an explicit generative model of the complete time course (Rousselet, 2025).

Finally, practical adoption also requires consideration of computational cost. It is important to note that the additional temporal precision and richer uncertainty quantification provided by the BGAM approach come at a computational cost. Conventional cluster-based permutation tests are highly optimised and remain substantially faster, making them particularly well suited for rapid exploratory analyses and large simulation studies. In the one-dimensional temporal case, cluster-based permutation tests completed in approximately 2 minutes per dataset (10,000 permutations), whereas the corresponding BGAM model required approximately 30 minutes (4 chains, 5,000 iterations each) on a recent laptop. Thus, BGAMs are not intended as a computationally equivalent replacement for permutation-based procedures. Rather, they trade additional computation time for a model-based estimate of the temporal structure of the effect, while accounting for group structures in the data and providing posterior evidence through time and direct estimates of effect onset and offset. Given that model fitting can be parallelised and that fitting time remains modest relative to the time invested in experimental design, data collection, preprocessing, and large-scale validation analyses, we view this computational cost as acceptable when the goal is accurate and robust temporal characterisation rather than only cluster-level significance testing (as noted by Spencer et al., 2022, in the context of spatial Bayesian GLMs applied to fMRI data).

To facilitate adoption, we developed the neurogam open-source R package, which implements the proposed method using brms. The package integrates seamlessly with MNE-Python (Gramfort, 2013), enabling researchers to process M/EEG data in Python and import them directly into R for model-based inference without cumbersome data export. This interoperability, described in Appendix D, is designed to encourage broader use of model-based approaches in cognitive neuroscience.

In conclusion, we introduced a model-based approach for temporally localised inference in M/EEG data that explicitly represents the temporal trajectory and between-participant variability. Across ground-truth simulations, the approach provided the most accurate overall recovery of the true temporal extent and comparatively accurate onset localisation across SNRs, although offset estimates were more variable and conventional procedures performed better on some offset-specific metrics. In empirical MEG data, BGAM also yielded the highest split-half temporal overlap and highly consistent temporal segmentation across repeated participant splits. These converging results support BGAMs as a flexible and useful complement to conventional procedures when explicit modelling of temporal structure, multilevel variation, and uncertainty in temporal localisation are scientifically important.

## Data and code availability

The simulation results as well as the R code to reproduce the simulations are available at: https://github.com/lnalborczyk/brms_meeg. The neurogam R package is available at https://github.com/lnalborczyk/neurogam.

## Packages

We used R v. 4.5.2 (R Core Team, 2025) and the following R packages: assertthat v. 0.2.1 (Wickham, 2019), brms v. 2.23.2 (Bürkner, 2017, 2018, 2021), ggpubr v. 0.6.3 (Kassambara, 2026), ggrepel v. 0.9.8 (Slowikowski, 2026), glue v. 1.8.1 (Hester & Bryan, 2026), grateful v. 0.3.0 (Rodriguez-Sanchez & Jackson, 2025), gt v. 1.3.0 (Iannone et al., 2026), knitr v. 1.52 (Xie, 2014, 2015, 2026), loo v. 2.10.1 (Magnusson et al., 2019a, 2019b; Paananen et al., 2021; Sivula et al., 2025; Vehtari et al., 2017, 2024a, 2026; Yao et al., 2018), MetBrewer v. 0.2.0 (Mills, 2022), pakret v. 0.3.1 (Gallou, 2026), patchwork v. 1.3.2 (T. L. Pedersen, 2025), posterior v. 1.7.0 (Bürkner et al., 2026; Lambert & Vehtari, 2022; Margossian et al., 2024; Vehtari et al., 2021b, 2024b), rmarkdown v. 2.32 (Allaire et al., 2026; Xie et al., 2018, 2020), scales v. 1.4.0 (Wickham et al., 2025), scico v. 1.5.0 (T. L. Pedersen & Crameri, 2023), tidybayes v. 3.0.7 (Kay, 2024), tidytext v. 0.4.3 (Silge & Robinson, 2016), tidyverse v. 2.0.0 (Wickham et al., 2019).

## Acknowledgements

We acknowledge the Centre de Calcul Intensif d’Aix-Marseille for granting access to its high performance computing resources.

## Appendix A Formal specification of the Bayesian additive multilevel model

Let *y_i_* denote the mean EEG signal for observation *i*, with known standard error *s_i_*, participant *j*[*i*], condition contrast *c_i_*, condition label *g_i_ ∈ {*1, 2*}*, and time *t_i_*. The model can be written as:

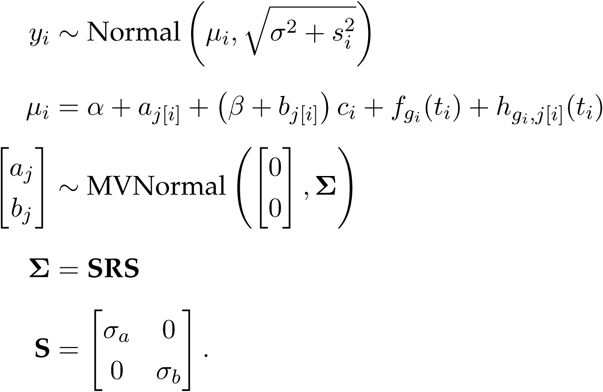

Here, *α* is the population-level intercept, *β* is the population-level condition effect, and *a_j_* and *b_j_* are participant-level deviations in intercept and condition effect (i.e., varying intercept and slope). The function *f_g_*(*t*) represents the population-level temporal trajectory for condition *g*, whereas *h_g,j_*(*t*) represents the participant-specific smooth deviation from that population trajectory. The residual scale combines the estimated residual standard deviation *σ* with the known standard error *s_i_* of each participant-level mean.

The population-level condition-specific smooths can be represented using basis functions as

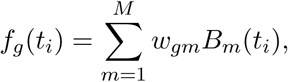

where *B_m_*(*t_i_*) are spline basis functions and *w_gm_* are condition-specific spline coefficients. Participant-specific smooth deviations can analogously be written as

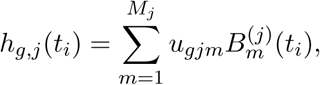

where *u_gjm_* are participant- and condition-specific spline coefficients. Their magnitude and smoothness are controlled by penalisation parameters estimated hierarchically from the data. The priors follow the default brms specification:

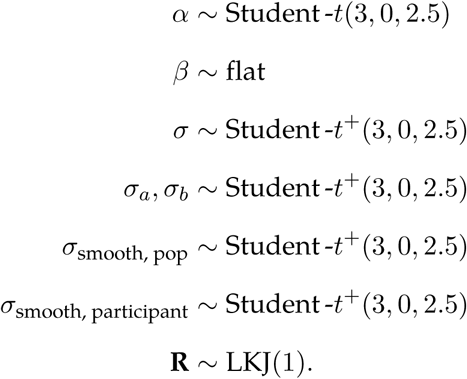

The superscript + indicates truncation to positive values. The correlation matrix **R** describes the correlation between participant-level intercepts and condition effects.

## Appendix B Application to 2D time-resolved decoding results (cross-temporal generalisation)

We conducted a cross-temporal generalisation analysis of the decoding data from Nalborczyk et al. (in preparation), in which we assessed the performance of classifiers trained and tested at various time points of the trial (King & Dehaene, 2014). This analysis was performed at the participant level, resulting in a 2D matrix where each element contains the decoding accuracy (ROC AUC) of a classifier trained at time point training*i* and tested at time point testing*j* for each participant (Figure B1).

**Figure B1.**
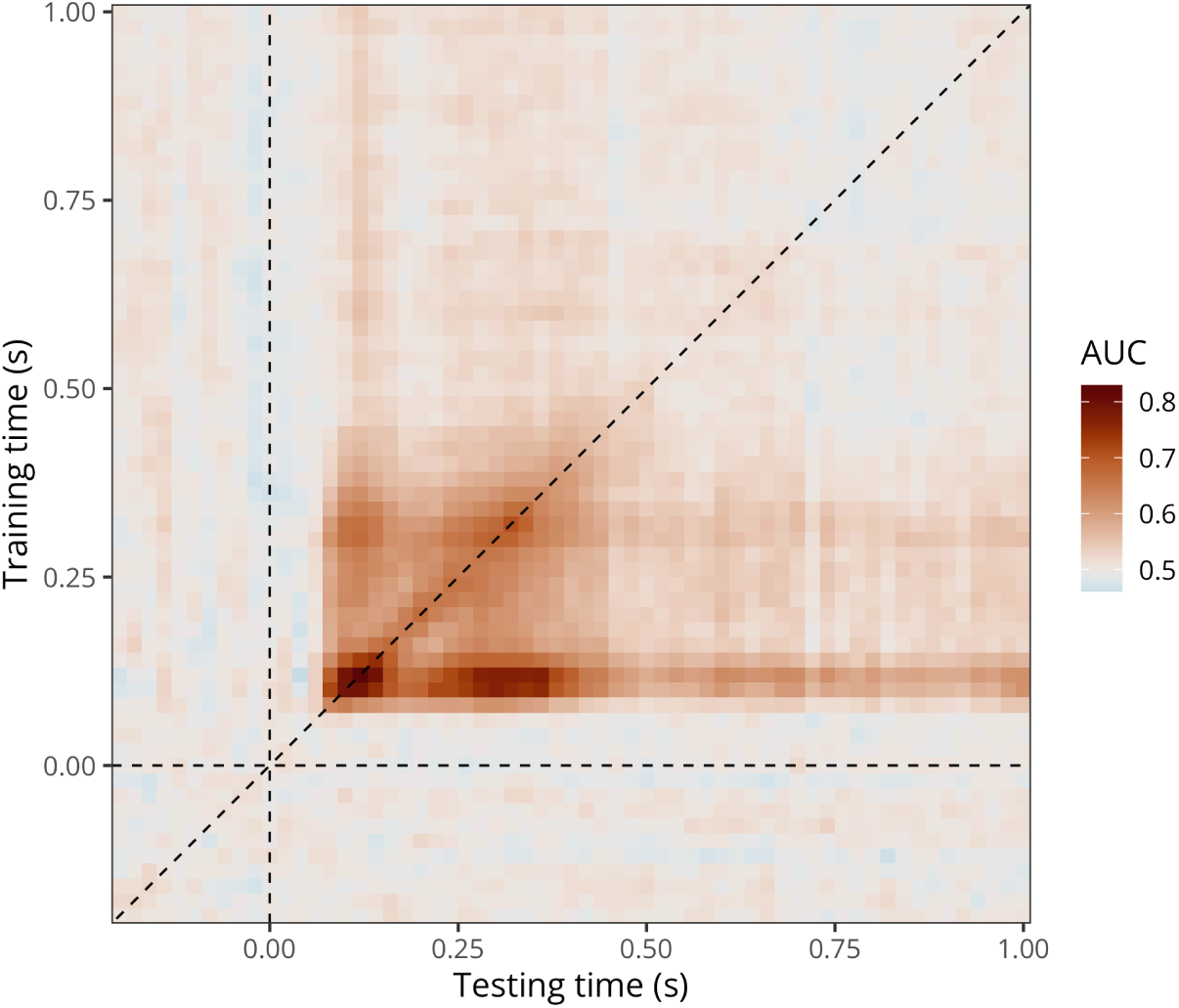
Group-level average cross-temporal generalisation matrix of decoding performance (data from Nalborczyk et al., in preparation).

To model cross-temporal generalisation matrices of decoding performance, we extended our initial BGAM to take into account the bivariate temporal distribution of AUC values, thus producing naturally smoothed estimates (time courses) of AUC values and posterior odds. This model can be written as follows:

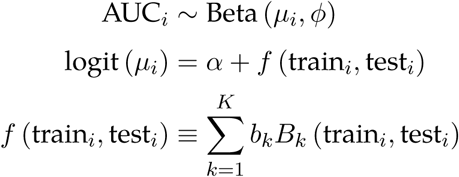

where AUC values are assumed to follow a Beta distribution, parametrised by a mean *µ_i_* and a precision parameter *ϕ*. The mean *µ_i_* is linked to the predictors through a logit link function. The smooth function *f* (train*_i_,* test*_i_*) represents a two-dimensional surface defined over training and testing times, which captures how decoding performance varies across the temporal generalisation matrix. This surface is approximated by a linear combination of *K* basis functions *B_k_*(*·, ·*), each weighted by a coefficient *b_k_*. The basis functions are constructed using a tensor product of univariate splines (here thin-plate splines) applied to the training and testing time dimensions (Wood, 2003, 2017).

We fitted this model using brms and the t2() tensor product smooth constructor with full penalties (E. J. Pedersen et al., 2019; Wood, 2017). We ran four MCMCs to approximate the posterior distribution, including each 3,000 iterations and a warmup of 1,000 iterations, yielding a total of 8,000 posterior samples to be used for inference.

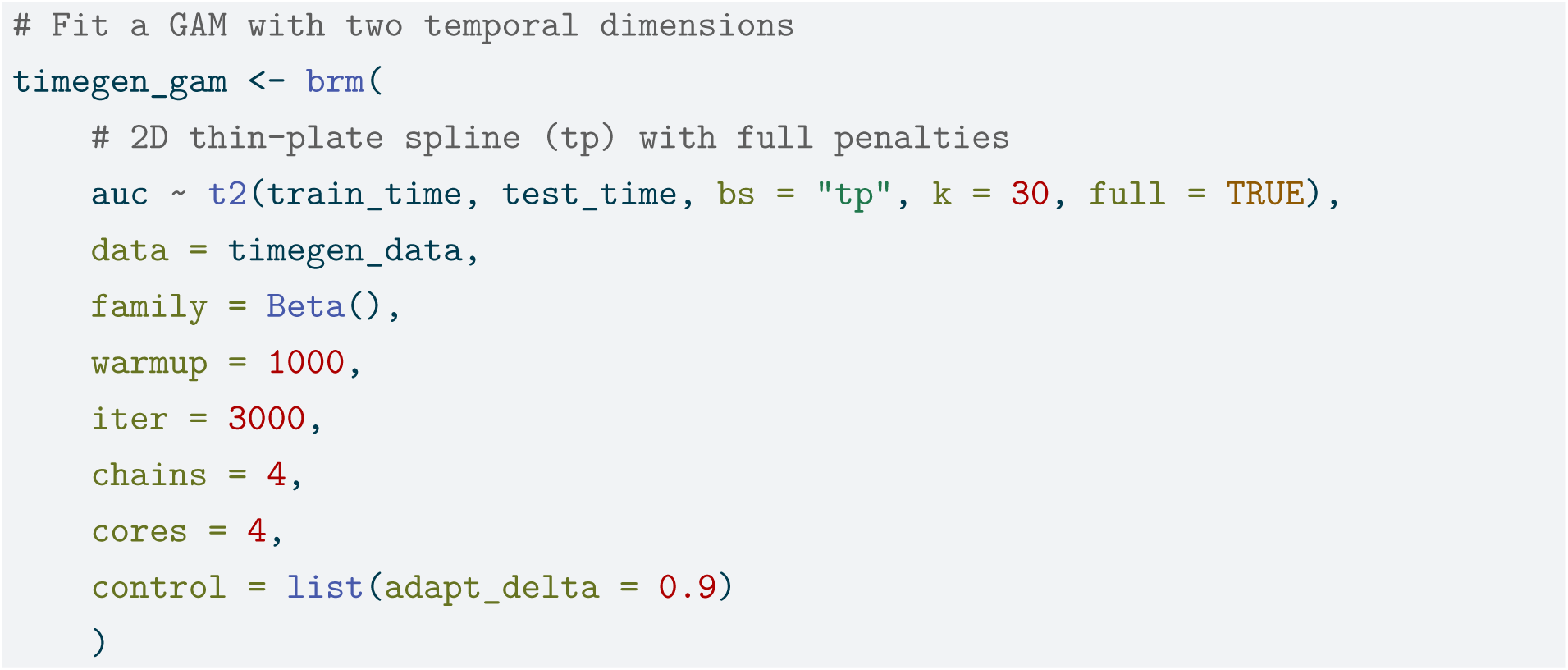

Figure B2 shows the model predictions (left) together with the region identified by thresholding the posterior odds (right). The model could be extended to a multilevel bivariate GAM by including participant-specific smooths, for example:

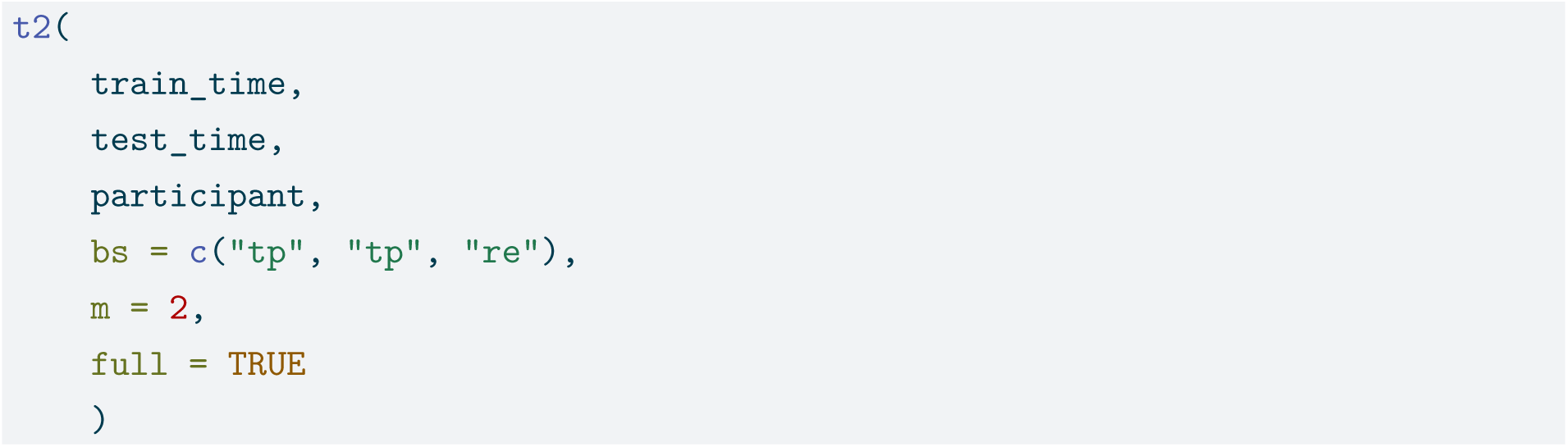

**Figure B2.**
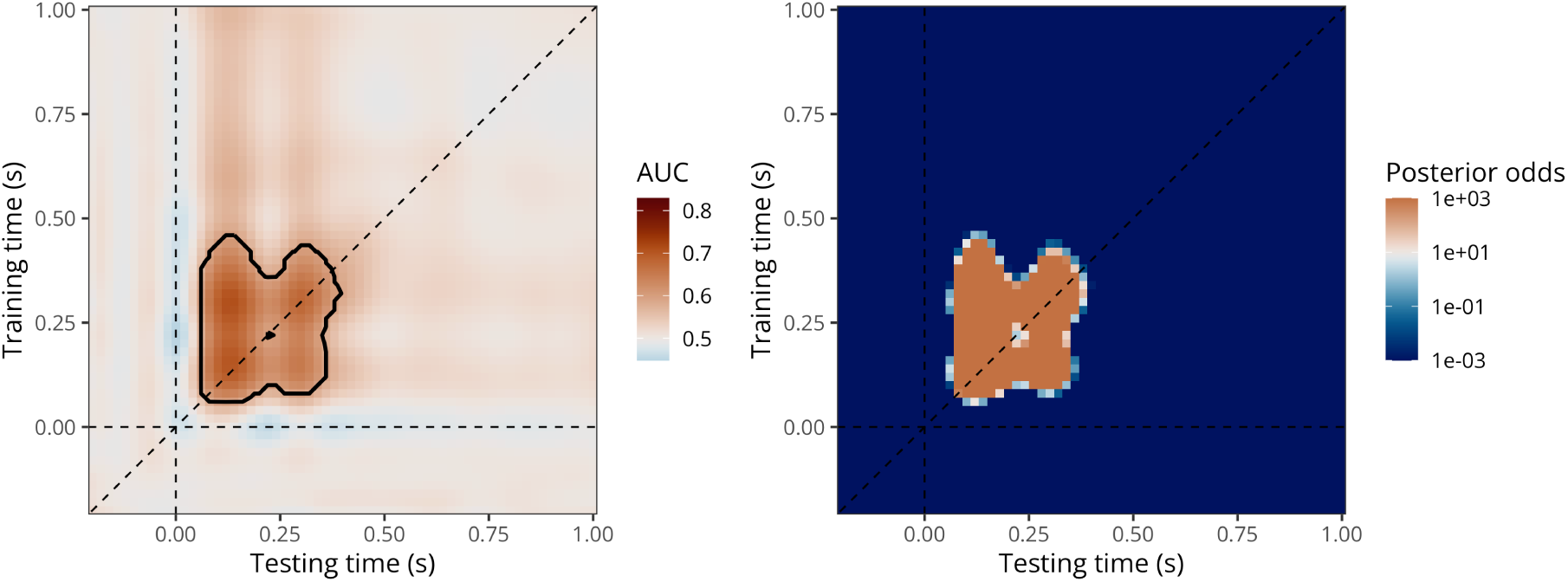
Predicted AUC values with threshold (left) and posterior odds of decoding accuracy being above chance (right) according to the bivariate BGAM.

**Table C1.**
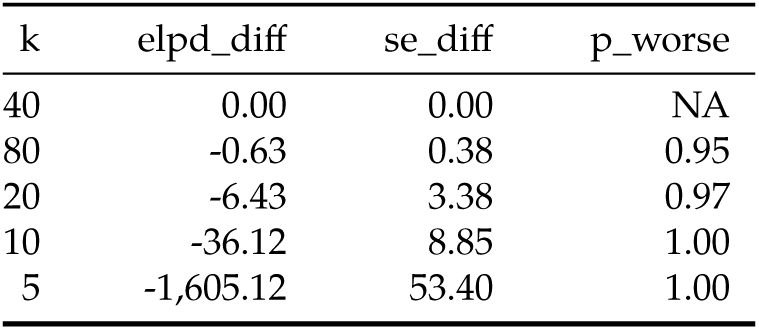
Model comparison using leave-one-out cross-validation. Models are arranged by the difference in expected log-pointwise density (ELPD) to the best model (i.e., k = 40).

## Appendix C How to choose the GAM basis dimension?

There is no universal recommendation for choosing the optimal value of k, as it depends on several factors, including the sampling rate, preprocessing steps (e.g., signal-to-noise ratio, low-pass filtering), and the underlying neural dynamics of the phenomenon under investigation. A practical strategy is to choose *k* sufficiently large that the basis dimension does not artificially constrain the smooth, and then assess its adequacy using posterior predictive checks and sensitivity analyses. Alternatively, models with different values of *k* can be compared using predictive criteria such as LOO. We illustrate this approach below.

Figure C1 presents the posterior predictions and two forms of posterior predictive checks (PPCs) for each GAM fit using different numbers of basis functions (*k ∈ {*5, 10, 20, 40, 80*}*). With the exception of the *k* = 5 model, all other fits yield satisfactory PPCs, indicating that the predicted data closely resemble the empirical observations. However, model comparison using the leave-one-out information criterion (LOOIC), as summarised in Table C1, identifies the *k* = 40 model as the best-performing one in terms of LOOIC, closely followed by the *k* = 80 model. Future simulation studies could further investigate how such model selection criteria relate to the precision of onset and offset estimates.

**Figure C1.**
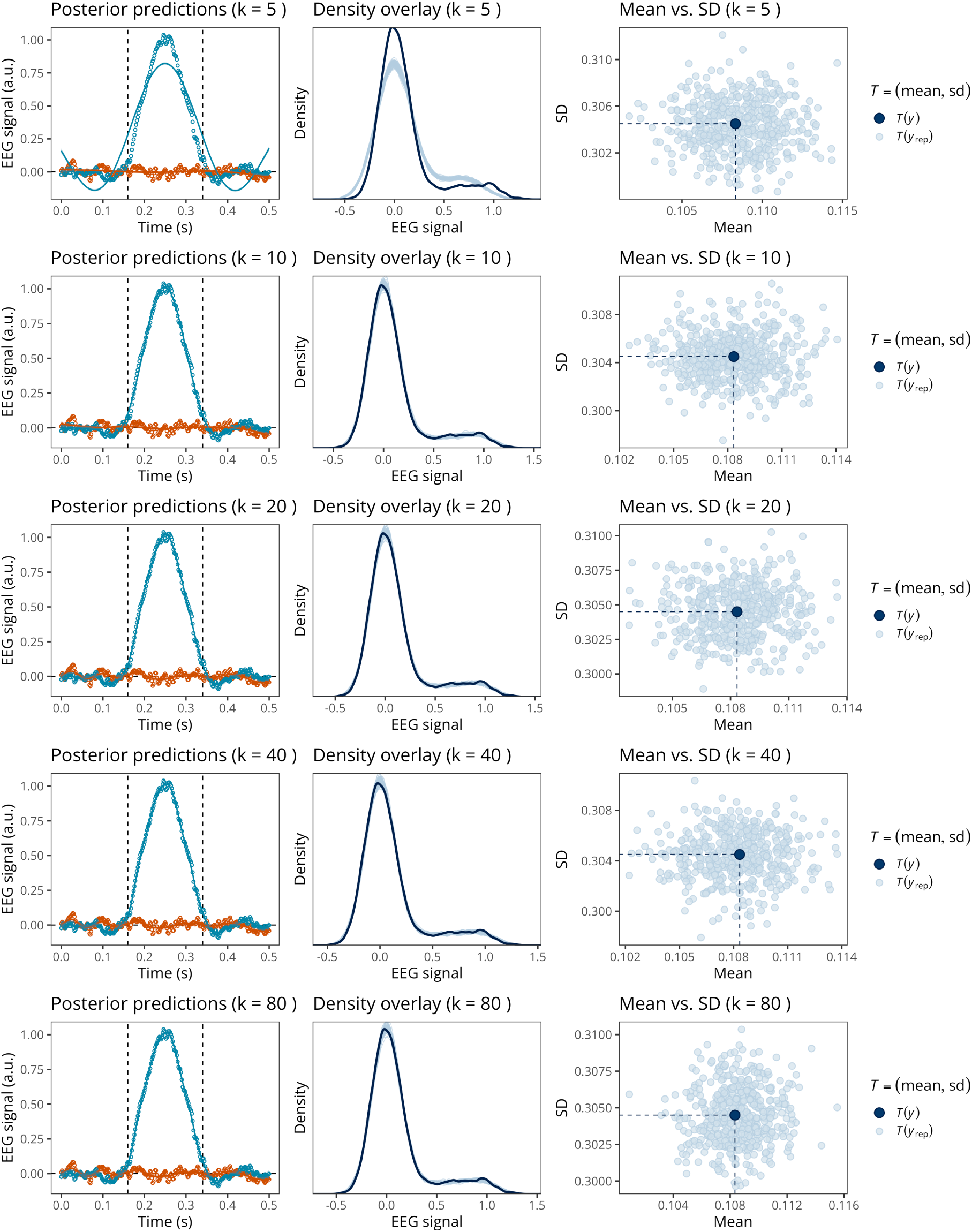
Posterior predictions and posterior predictive checks for the GAM with varying k (in rows).

## Appendix D R package and integration with MNE-Python

For readers who are already familiar with brms, the recommended pipeline is to use the code provided in the main paper (available at https://github.com/lnalborczyk/brms_meeg). It is also possible to call functions from the neurogam R package (available at https://github.com/ lnalborczyk/neurogam) which come with sensible defaults.

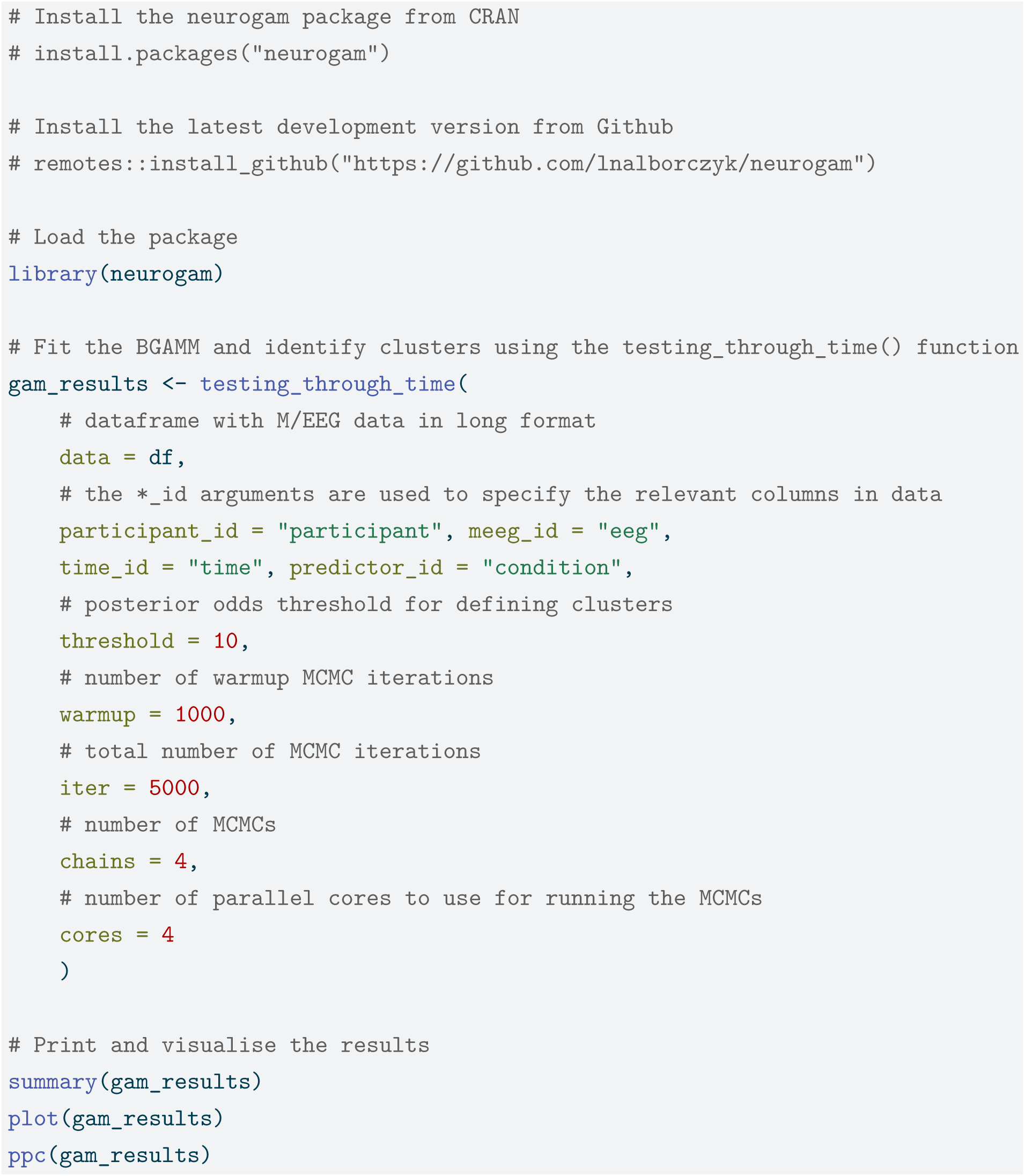

The neurogam package can also be called from Python using the rpy2 module, and can easily be integrated into MNE-Python pipelines. For example, we use it below to estimate the onset and offset of effects for one EEG channel from an MNE evoked object. The code used to reshape the sample MNE dataset is available in the online supplementary materials, and we further refer to the MNE documentation about converting MNE epochs to Pandas dataframes in long format (i.e., with one observation per row).

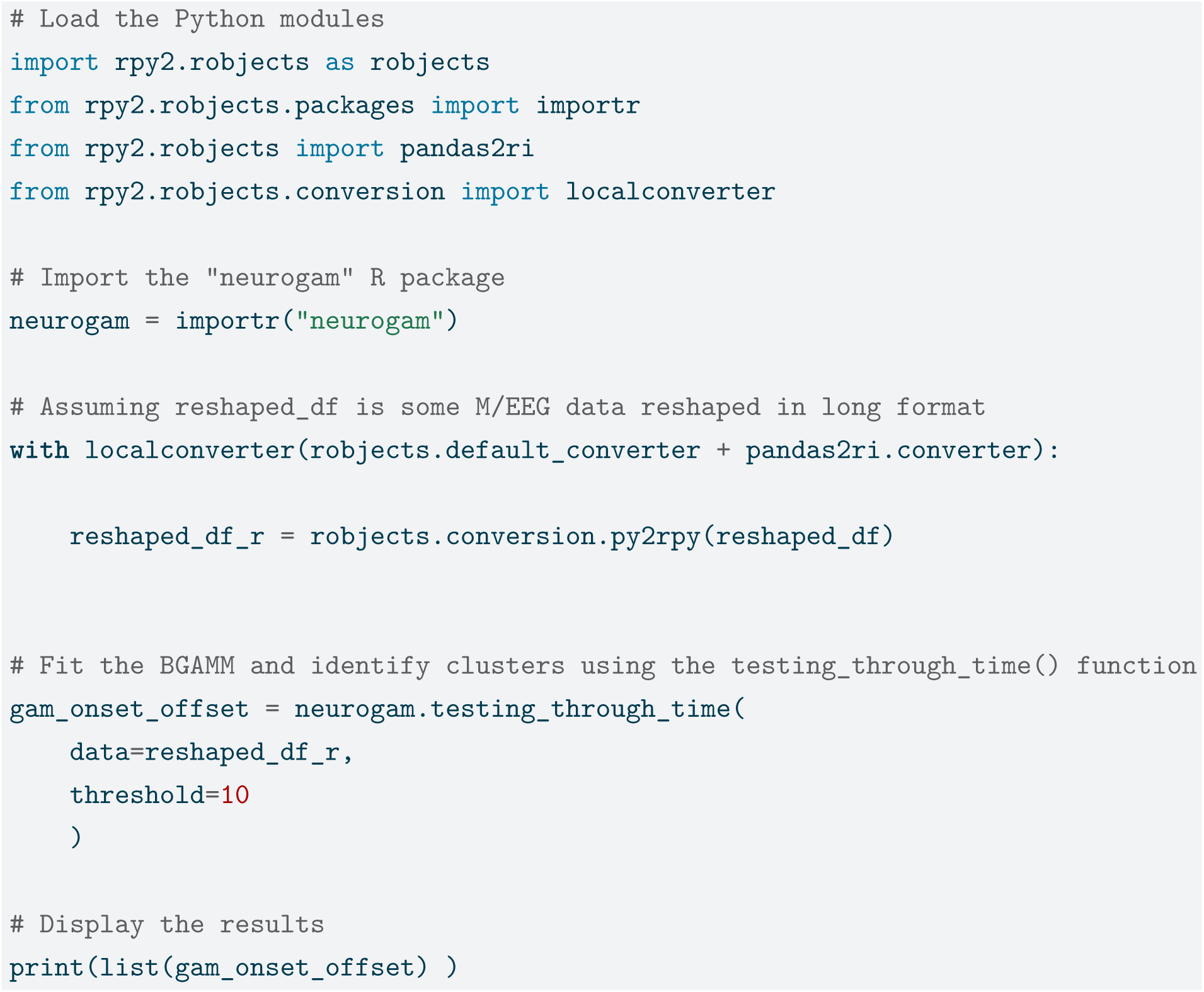

Similarly, neurogam can be called from Julia via the RCall.jl interface, allowing users to run the same R functions from Julia workflows without leaving the Julia environment.

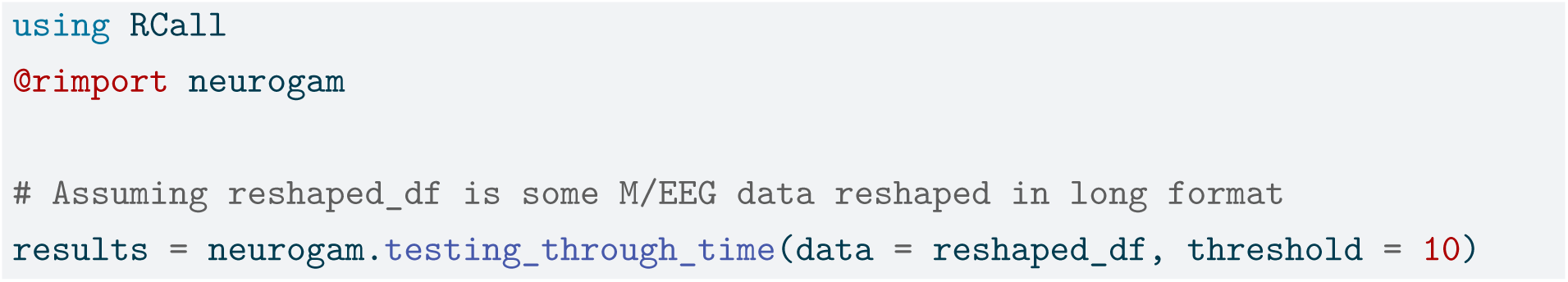

## Notes

https://github.com/lnalborczyk/brms_meeg

### Competing Interest Statement

The authors have declared no competing interest.

### Summary of Updates

Updated simulation study and methods description.

## References

Abugaber, D., Finestrat, I., Luque, A., & Morgan-Short, K. (2023). Generalized additive mixed modeling of EEG supports dual-route accounts of morphosyntax in suggesting no word frequency effects on processing of regular grammatical forms. Journal of Neurolinguistics, 67, 101137. 10.1016/j.jneuroling.2023.101137

Allaire, J., Xie, Y., Dervieux, C., McPherson, J., Luraschi, J., Ushey, K., Atkins, A., Wickham, H., Cheng, J., Chang, W., & Iannone, R. (2026). rmarkdown: Dynamic documents for r. https://github.com/rstudio/rmarkdown

Baayen, R. H., & Linke, M. (2020). Generalized Additive Mixed Models (pp. 563–591). Springer International Publishing. 10.1007/978-3-030-46216-1_23

Baayen, R. H., Rij, J. van, Cat, C. de, & Wood, S. (2018). Autocorrelated errors in experimental data in the language sciences: Some solutions offered by generalized additive mixed models (pp. 49–69). Springer International Publishing. 10.1007/978-3-319-69830-4_4

Benjamini, Y., & Hochberg, Y. (1995). Controlling the False Discovery Rate: A Practical and Powerful Approach to Multiple Testing. Journal of the Royal Statistical Society Series B: Statistical Methodology, 57(1), 289–300. 10.1111/j.2517-6161.1995.tb02031.x

Benjamini, Y., & Yekutieli, D. (2001). The control of the false discovery rate in multiple testing under dependency. The Annals of Statistics, 29(4). 10.1214/aos/1013699998

Bullmore, E. T., Suckling, J., Overmeyer, S., Rabe-Hesketh, S., Taylor, E., & Brammer, M. J. (1999). Global, voxel, and cluster tests, by theory and permutation, for a difference between two groups of structural MR images of the brain. IEEE Transactions on Medical Imaging, 18(1), 32–42. 10.1109/42.750253

Bürkner, P.-C. (2017). brms: An R package for Bayesian multilevel models using Stan. Journal of Statistical Software, 80(1), 1–28. 10.18637/jss.v080.i01

Bürkner, P.-C. (2018). Advanced Bayesian multilevel modeling with the R package brms. The R Journal, 10(1), 395–411. 10.32614/RJ-2018-017

Bürkner, P.-C. (2021). Bayesian item response modeling in R with brms and Stan. Journal of Statistical Software, 100(5), 1–54. 10.18637/jss.v100.i05

Bürkner, P.-C., Gabry, J., Kay, M., & Vehtari, A. (2026). posterior: Tools for working with posterior distributions. https://mc-stan.org/posterior/

Delorme, A., & Makeig, S. (2004). EEGLAB: an open source toolbox for analysis of single-trial EEG dynamics including independent component analysis. Journal of Neuroscience Methods, 134(1), 9–21. 10.1016/j.jneumeth.2003.10.009

Dinga, R., Fraza, C. J., Bayer, J. M. M., Kia, S. M., Beckmann, C. F., & Marquand, A. F. (2021). Normative modeling of neuroimaging data using generalized additive models of location scale and shape. 10.1101/2021.06.14.448106

Dunagan, D., Jordan, T., Hale, J. T., Pylkkänen, L., & Chacón, D. A. (2025). Evaluating the timecourses of morpho-orthographic, lexical, and grammatical processing following rapid parallel visual presentation: An EEG investigation in English. Cognition, 257, 106080. 10.1016/j.cognition.2025.106080

Ehinger, B. V., & Dimigen, O. (2019). Unfold: An integrated toolbox for overlap correction, non-linear modeling, and regression-based EEG analysis. PeerJ, 7, e7838. 10.7717/peerj.7838

Frossard, J., & Renaud, O. (2022). The cluster depth tests: Toward point-wise strong control of the family-wise error rate in massively univariate tests with application to M/EEG. NeuroImage, 247, 118824. 10.1016/j.neuroimage.2021.118824

Gabry, J., Simpson, D., Vehtari, A., Betancourt, M., & Gelman, A. (2019). Visualization in Bayesian workflow. Journal of the Royal Statistical Society: Series A (Statistics in Society), 182(2), 389–402. 10.1111/rssa.12378

Gallou, A. (2026). pakret: Cite “R” packages on the fly in “R Markdown” and “Quarto”. 10.32614/CRAN.package.pakret

Gelman, A., Vehtari, A., Simpson, D., Margossian, C. C., Carpenter, B., Yao, Y., Kennedy, L., Gabry, J., Bürkner, P.-C., & Modrák, M. (2020). Bayesian workflow. arXiv:2011.01808 [Stat]. http://arxiv.org/abs/2011.01808

Gramfort, A. (2013). MEG and EEG data analysis with MNE-python. Frontiers in Neuroscience, 7. 10.3389/fnins.2013.00267

Hastie, T. J., & Tibshirani, R. J. (2017). Generalized Additive Models. Routledge. 10.1201/9780203753781

Hester, J., & Bryan, J. (2026). glue: Interpreted string literals. 10.32614/CRAN.package.glue

Holm, S. (1979). A simple sequentially rejective multiple test procedure. Scandinavian Journal of Statistics, 6(2), 65–70. http://www.jstor.org/stable/4615733

Iannone, R., Cheng, J., Schloerke, B., Haughton, S., Hughes, E., Lauer, A., François, R., Seo, J., Brevoort, K., & Roy, O. (2026). gt: Easily create presentation-ready display tables. 10.32614/CRAN.package.gt

Kassambara, A. (2026). ggpubr: “ggplot2” based publication ready plots. 10.32614/CRAN.package.ggpubr

Kay, M. (2024). tidybayes: Tidy data and geoms for Bayesian models. 10.5281/zenodo.1308151

Killick, R., Haynes, K., & Eckley, I. A. (2022). changepoint: An R package for changepoint analysis. https://CRAN.R-project.org/package=changepoint

King, J.-R., & Dehaene, S. (2014). Characterizing the dynamics of mental representations: the temporal generalization method. Trends in Cognitive Sciences, 18(4), 203–210. 10.1016/j.tics.2014.01.002

Kruschke, J. K., & Liddell, T. M. (2017). The Bayesian New Statistics: Hypothesis testing, estimation, meta-analysis, and power analysis from a Bayesian perspective. Psychonomic Bulletin & Review, 25(1), 178–206. 10.3758/s13423-016-1221-4

Lambert, B., & Vehtari, A. (2022). Rstar: A robust MCMC convergence diagnostic with uncertainty using decision tree classifiers. Bayesian Analysis, 17(2), 353–379. 10.1214/20-BA1252

Magnusson, M., Andersen, M. R., Jonasson, J., & Vehtari, A. (2019a). Leave-one-out cross-validation for large data. Thirty-Sixth International Conference on Machine Learning, 97, 4244–4253.

Magnusson, M., Andersen, M. R., Jonasson, J., & Vehtari, A. (2019b). Leave-one-out cross-validation for model comparison in large data. Proceedings of the 23rd International Conference on Artificial Intelligence and Statistics (AISTATS), 108, 341–351.

Margossian, C. C., Hoffman, M. D., Sountsov, P., Riou-Durand, L., Vehtari, A., & Gelman, A. (2024). Nested rhat: Assessing the convergence of markov chain monte carlo when running many short chains. Bayesian Analysis. 10.1214/24-BA1453

Maris, E. (2011). Statistical testing in electrophysiological studies. Psychophysiology, 49(4), 549–565. 10.1111/j.1469-8986.2011.01320.x

Maris, E., & Oostenveld, R. (2007). Nonparametric statistical testing of EEG- and MEG-data. Journal of Neuroscience Methods, 164(1), 177–190. 10.1016/j.jneumeth.2007.03.024

Meulman, N., Sprenger, S. A., Schmid, M. S., & Wieling, M. (2023). GAM-based individual difference measures for L2 ERP studies. Research Methods in Applied Linguistics, 2(3), 100079. 10.1016/j.rmal.2023.100079

Meulman, N., Wieling, M., Sprenger, S. A., Stowe, L. A., & Schmid, M. S. (2015). Age Effects in L2 Grammar Processing as Revealed by ERPs and How (Not) to Study Them. PLOS ONE, 10(12), e0143328. 10.1371/journal.pone.0143328

Miller, D. L. (2025). Bayesian views of generalized additive modelling. Methods in Ecology and Evolution. 10.1111/2041-210x.14498

Mills, B. R. (2022). MetBrewer: Color palettes inspired by works at the metropolitan museum of art. 10.32614/CRAN.package.MetBrewer

Nalborczyk, L., Batailler, C., Lœvenbruck, H., Vilain, A., & Bürkner, P.-C. (2019). An Introduction to Bayesian Multilevel Models Using brms: A Case Study of Gender Effects on Vowel Variability in Standard Indonesian. Journal of Speech, Language, and Hearing Research, 62(5), 1225–1242. 10.1044/2018_jslhr-s-18-0006

Nalborczyk, L., Hauw, F., Torcy, H. de, Dehaene, S., & Cohen, L. (in preparation). Neural and representational dynamics of tickertape synesthesia.

Paananen, T., Piironen, J., Buerkner, P.-C., & Vehtari, A. (2021). Implicitly adaptive importance sampling. Statistics and Computing, 31, 16.

Pedersen, E. J., Miller, D. L., Simpson, G. L., & Ross, N. (2019). Hierarchical generalized additive models in ecology: An introduction with mgcv. PeerJ, 7, e6876. 10.7717/peerj.6876

Pedersen, T. L. (2025). patchwork: The composer of plots. 10.32614/CRAN.package.patchwork

Pedersen, T. L., & Crameri, F. (2023). scico: Colour palettes based on the scientific colour-maps. 10.32614/CRAN.package.scico

Pernet, C. R., Chauveau, N., Gaspar, C., & Rousselet, G. A. (2011). LIMO EEG: A Toolbox for Hierarchical LInear MOdeling of ElectroEncephaloGraphic Data. Computational Intelligence and Neuroscience, 2011, 1–11. 10.1155/2011/831409

Pernet, C. R., Latinus, M., Nichols, T. E., & Rousselet, G. A. (2015). Cluster-based computational methods for mass univariate analyses of event-related brain potentials/fields: A simulation study. Journal of Neuroscience Methods, 250, 85–93. 10.1016/j.jneumeth.2014.08.003

R Core Team. (2025). R: A language and environment for statistical computing. R Foundation for Statistical Computing. https://www.R-project.org/

Rasmussen, C. E., & Williams, C. K. I. (2005). Gaussian Processes for Machine Learning. 10.7551/mitpress/3206.001.0001

Rigby, R. A., & Stasinopoulos, D. M. (2005). Generalized Additive Models for Location, Scale and Shape. Journal of the Royal Statistical Society Series C: Applied Statistics, 54(3), 507–554. 10.1111/j.1467-9876.2005.00510.x

Rij, J. van, Hendriks, P., Rijn, H. van, Baayen, R. H., & Wood, S. N. (2019). Analyzing the Time Course of Pupillometric Data. Trends in Hearing, 23. 10.1177/2331216519832483

Riutort-Mayol, G., Bürkner, P.-C., Andersen, M. R., Solin, A., & Vehtari, A. (2023). Practical Hilbert space approximate Bayesian Gaussian processes for probabilistic programming. Statistics and Computing, 33(1), 17. 10.1007/s11222-022-10167-2

Rodriguez-Sanchez, F., & Jackson, C. P. (2025). grateful: Facilitate citation of R packages. https://pakillo.github.io/grateful/

Rousselet, G. A. (2025). Using cluster-based permutation tests to estimate MEG/EEG onsets: How bad is it? European Journal of Neuroscience, 61(1), e16618. 10.1111/ejn.16618

Rousselet, G. A., Pernet, C. R., Bennett, P. J., & Sekuler, A. B. (2008). Parametric study of EEG sensitivity to phase noise during face processing. BMC Neuroscience, 9(1). 10.1186/1471-2202-9-98

Sassenhagen, J., & Draschkow, D. (2019). Cluster-based permutation tests of MEG/EEG data do not establish significance of effect latency or location. Psychophysiology, 56(6). 10.1111/psyp.13335

Silge, J., & Robinson, D. (2016). tidytext: Text mining and analysis using tidy data principles in r. JOSS, 1(3). 10.21105/joss.00037

Sivula, T., Magnusson, M., Matamoros, A. A., & Vehtari, A. (2025). Uncertainty in bayesian leave-one-out cross-validation based model comparison. Bayesian Analysis.

Skukies, R., & Ehinger, B. (2021). Modelling event duration and overlap during EEG analysis. Journal of Vision, 21(9), 2037. 10.1167/jov.21.9.2037

Skukies, R., Schepers, J., & Ehinger, B. (2024, December 9). Brain responses vary in duration - modeling strategies and challenges. 10.1101/2024.12.05.626938

Slowikowski, K. (2026). ggrepel: Automatically position non-overlapping text labels with “ggplot2”. 10.32614/CRAN.package.ggrepel

Smith, N. J., & Kutas, M. (2014a). Regression-based estimation of ERP waveforms: I. The rERP framework. Psychophysiology, 52(2), 157–168. 10.1111/psyp.12317

Smith, N. J., & Kutas, M. (2014b). Regression-based estimation of ERP waveforms: II. Nonlinear effects, overlap correction, and practical considerations. Psychophysiology, 52(2), 169–181. 10.1111/psyp.12320

Smith, S., & Nichols, T. (2009). Threshold-free cluster enhancement: Addressing problems of smoothing, threshold dependence and localisation in cluster inference. NeuroImage, 44(1), 83–98. 10.1016/j.neuroimage.2008.03.061

Sóskuthy, M. (2021). Evaluating generalised additive mixed modelling strategies for dynamic speech analysis. Journal of Phonetics, 84, 101017. 10.1016/j.wocn.2020.101017

Spencer, D., Yue, Y. R., Bolin, D., Ryan, S., & Mejia, A. F. (2022). Spatial bayesian GLM on the cortical surface produces reliable task activations in individuals and groups. NeuroImage, 249, 118908. 10.1016/j.neuroimage.2022.118908

Tremblay, A., & Newman, A. J. (2014). Modeling nonlinear relationships in ERP data using mixed-effects regression with R examples. Psychophysiology, 52(1), 124–139. 10.1111/psyp.12299

Umlauf, N., Klein, N., & Zeileis, A. (2018). BAMLSS: Bayesian Additive Models for Location, Scale, and Shape (and Beyond). Journal of Computational and Graphical Statistics, 27(3), 612–627. 10.1080/10618600.2017.1407325

Ushey, K., Allaire, J., & Tang, Y. (2024). Reticulate: Interface to ‘python’. https://CRAN.R-project.org/package=reticulate

Vehtari, A., Gabry, J., Magnusson, M., Yao, Y., Bürkner, P.-C., Paananen, T., & Gelman, A. (2026). loo: Efficient leave-one-out cross-validation and WAIC for bayesian models. https://mc-stan.org/loo/

Vehtari, A., Gelman, A., & Gabry, J. (2017). Practical bayesian model evaluation using leave- one-out cross-validation and WAIC. Statistics and Computing, 27, 1413–1432. 10.1007/s11222-016-9696-4

Vehtari, A., Gelman, A., Simpson, D., Carpenter, B., & Bürkner, P.-C. (2021a). Rank-normalization, folding, and localization: An improved R^ for assessing convergence of MCMC (with discussion). Bayesian Analysis, 16(2). 10.1214/20-ba1221

Vehtari, A., Gelman, A., Simpson, D., Carpenter, B., & Bürkner, P.-C. (2021b). Rank-normalization, folding, and localization: An improved rhat for assessing convergence of MCMC (with discussion). Bayesian Analysis, 16(2), 667–718.

Vehtari, A., Simpson, D., Gelman, A., Yao, Y., & Gabry, J. (2024a). Pareto smoothed importance sampling. Journal of Machine Learning Research, 25(72), 1–58.

Vehtari, A., Simpson, D., Gelman, A., Yao, Y., & Gabry, J. (2024b). Pareto smoothed importance sampling. Journal of Machine Learning Research, 25(72), 1–58.

Wickham, H. (2019). assertthat: Easy pre and post assertions. 10.32614/CRAN.package.assertthat

Wickham, H., Averick, M., Bryan, J., Chang, W., McGowan, L. D., François, R., Grolemund, G., Hayes, A., Henry, L., Hester, J., Kuhn, M., Pedersen, T. L., Miller, E., Bache, S. M., Müller, K., Ooms, J., Robinson, D., Seidel, D. P., Spinu, V., … Yutani, H. (2019). Welcome to the tidyverse. Journal of Open Source Software, 4(43), 1686. 10.21105/joss.01686

Wickham, H., Pedersen, T. L., & Seidel, D. (2025). scales: Scale functions for visualization. 10.32614/CRAN.package.scales

Wieling, M. (2018). Analyzing dynamic phonetic data using generalized additive mixed modeling: A tutorial focusing on articulatory differences between L1 and L2 speakers of English. Journal of Phonetics, 70, 86–116. 10.1016/j.wocn.2018.03.002

Wood, S. N. (2003). Thin Plate Regression Splines. Journal of the Royal Statistical Society Series B: Statistical Methodology, 65(1), 95–114. 10.1111/1467-9868.00374

Wood, S. N. (2017). Generalized Additive Models. Chapman; Hall/CRC. 10.1201/9781315370279

Wüllhorst, V., Wüllhorst, R., Overmeyer, R., & Endrass, T. (2025). Comprehensive Analysis of Event-Related Potentials of Response Inhibition: The Role of Negative Urgency and Compulsivity. Psychophysiology, 62(2). 10.1111/psyp.70000

Xie, Y. (2014). knitr: A comprehensive tool for reproducible research in R. In V. Stodden, F. Leisch, & R. D. Peng (Eds.), Implementing reproducible computational research. Chapman; Hall/CRC.

Xie, Y. (2015). Dynamic documents with R and knitr (2nd ed.). Chapman; Hall/CRC. https://yihui.org/knitr/

Xie, Y. (2026). knitr: A general-purpose package for dynamic report generation in R. https://yihui.org/knitr/

Xie, Y., Allaire, J. J., & Grolemund, G. (2018). R markdown: The definitive guide. Chapman; Hall/CRC. https://pkg.yihui.org/rmarkdown-book/

Xie, Y., Dervieux, C., & Riederer, E. (2020). R markdown cookbook. Chapman; Hall/CRC. https://pkg.yihui.org/rmarkdown-cookbook/

Yao, Y., Vehtari, A., Simpson, D., & Gelman, A. (2018). Using stacking to average bayesian predictive distributions. Bayesian Analysis, 13, 917–1007. 10.1214/17-BA1091

